# Anatomically matched spatial comparisons identify *Trps1* as a regulator of knee interzone formation

**DOI:** 10.64898/2026.10.28.762640

**Authors:** Ji-Hye Yea, Sharon E.S. Onggo, Haneul Cho, Patrick Cahan

## Abstract

How the interzone forms and gives rise to articular cartilage, ligaments, and synovium is incompletely understood. Using our previously generated spatial and single-cell transcriptomic atlas of the mouse hindlimb, we compared the knee interzone with neighboring tissues and with the ankle interzone at matched positions, and identified *Trps1* as a knee interzone-enriched candidate. Pseudotime analysis placed *Trps1*-high cells in a transitional state between chondrogenic progenitors and matrix-producing chondrocytes, with higher chondrogenic and lower fibroblastic program scores. Targeted *Trps1* knockdown in hindlimb explants disrupted joint formation, permitted *Col2a1* expression within the interzone, and reduced adjacent cartilage formation. The associated chondrogenic program was conserved in the human embryonic knee, whereas *TRPS1* expression in osteoarthritic cartilage was associated with a fibroblastic matrix program. These results support a role for *Trps1* in knee interzone development and show that anatomically matched spatial comparison with explant perturbation can nominate and test joint regulators.

## Introduction

Synovial joints form at the interzone, a transient population of flattened, densely packed mesenchymal cells that emerges between developing skeletal elements and contributes to articular cartilage, intra-articular ligaments, and the synovial lining ^1–3^. Joint formation requires the spatial organization and differentiation of these populations alongside the developing cartilage templates. Disruption of this process can cause defects ranging from abnormal joint architecture to skeletal fusion ^4,5^. Altered joint morphology can also influence susceptibility to osteoarthritis (OA) later in life ^6,7^. Identifying regulators of early joint formation is therefore important for understanding both joint development and the developmental contributions to joint disease.

Genetic and lineage-tracing studies have established several features of interzone formation. *Gdf5* marks early joint progenitors, and combined loss of *Gdf5* and *Gdf6* causes joint fusions and skeletal patterning defects ^4,5^. *Wnt9a* promotes joint formation and suppresses chondrogenic differentiation within the interzone, while *Creb5* regulates signaling required for joint formation and articular cartilage development ^8–10^. Lineage tracing shows that *Gdf5*-expressing cells contribute to multiple joint tissues and enter the interzone over several days, with their time of arrival associated with their eventual contribution to joint structures^11,12^. Our previous single-cell study further revealed transcriptional heterogeneity among *Gdf5*-lineage progenitors at E12.5, before a morphologically distinct interzone becomes apparent ^13,14^. These findings establish the diversity and dynamic recruitment of joint progenitors, but the regulators that contribute to their differentiation and organization during early knee joint formation remain incompletely understood.

Single-cell RNA sequencing provides a basis for identifying population-associated genes and regulatory programs, but tissue dissociation removes the positional information needed to relate these profiles directly to the developing joint. A gene enriched in a transcriptionally defined population may also be expressed in related cells outside the prospective interzone. Moreover, cell identities are commonly assigned using established markers, as illustrated by single-cell studies of limb and joint development ^15,16^. Although such annotation enables investigation of population-associated pathways, it does not by itself establish enrichment at a particular anatomical site and developmental stage. Spatial transcriptomics complements single-cell profiling by retaining gene expression within its tissue context. Comparisons of the interzone with flanking cartilage and surrounding mesenchyme across developmental stages, followed by comparisons between knee and ankle interzones, can therefore prioritize knee interzone-enriched candidates without preselecting genes based on known pathways or regulatory functions. Spatial enrichment alone, however, does not establish a gene’s functional contribution to joint development. Functional testing must also account for the tissue architecture and interactions within which joint formation occurs. The time and resources required to generate conditional mouse models also limit the initial functional evaluation of newly identified candidates ^17,18^. Ex vivo limb explant culture preserves the anatomical relationships among developing tissues and supports continued skeletal development ^19,20^. Combined with localized siRNA delivery, this system allows candidate genes to be perturbed at the prospective joint while monitoring the development of the interzone and neighboring structures ^21^. Linking spatial candidate identification to functional perturbation thus provides an approach to investigating regulators of knee joint formation and their contributions to chondrogenesis.

Developmental regulators may also be relevant to changes in adult cartilage. A genome-wide association meta-analysis identified 77 high-confidence OA effector genes, of which 63 were associated with skeletal development, suggesting links between OA susceptibility, the establishment of joint structure, and responses to joint damage ^6^. Articular chondrocytes in OA can express a hypertrophic program resembling that of growth plate chondrocytes, while single-cell analyses identify fibrochondrocyte states associated with cartilage degeneration ^22–25^. These observations raise the question of whether regulators associated with embryonic joint formation retain similar transcriptional associations in adult cartilage or become associated with different programs in disease. Here, we combine spatial and single-cell transcriptomics of the mouse hindlimb at E11.5– E14.5 with anatomically matched comparisons across tissues, developmental stages, and joints to identify *Trps1* as a knee interzone-enriched candidate regulator. We characterize its expression across joint-associated populations and examine its relationship to chondrogenic differentiation. We then use localized knockdown in hindlimb explants to test its contribution to knee joint formation and chondrogenesis. Finally, we investigate whether the associated expression patterns and transcriptional programs are conserved in human embryonic development and altered in osteoarthritic cartilage.

## Results

### Anatomically matched spatial comparisons nominate *Trps1* as an interzone-enriched candidate regulator

We asked whether comparing the interzone with its neighboring tissues at matched positions in two joints would nominate interzone regulators. For this we drew on our recently released spatial and single-cell transcriptomic atlas of the mouse hindlimb, which profiled *Sox9*-IRES-GFP embryos from E11.5 to E15.5 and annotated cell populations by their position in the tissue ^26^. We used its E11.5 to E14.5 spatial sections, 43 sections containing 1,268,878 spots, together with histology and scRNA-seq of the same stages (Figure 1A). Sox9-GFP fluorescence and Safranin O/Fast Green (Saf-O/FG) staining outlined the femur and tibia cartilage anlagen and showed a *Sox9*-GFP-low cleft between them at approximately E13.5, the morphological onset of joint formation (Figure 1B). Within these sections, the cartilage markers *Col2a1* and *Sox9* and the interzone marker *Gdf5* distinguished cartilage from interzone at each stage (Figure 1C, D and Figure S1). To nominate candidates, we relied on a combination of expression-detection, temporal-enrichment, tissue-enrichment, and inter-joint comparisons. We first used Col2a1 and Sox9 expression together with tissue morphology to annotate anatomically matched regions—the interzone, central cartilage anlagen, and surrounding non-cartilage tissue— independently for the knee (femur–tibia) and ankle (tibia–talus) (Figure 1E). We retained genes detected in at least 5% of knee interzone spots pooled across E12.5 and E13.5, yielding 1,616 genes. Second, we retained genes enriched in the pooled E12.5–E13.5 knee interzone relative to both the E11.5 cartilage primordium and the E14.5 knee interzone, yielding 66 genes (Figure 1F). In a separate analysis of all genes, we compared the knee interzone with surrounding non-interzone tissues separately at E12.5 and E13.5. Twenty-five genes were enriched in the interzone at both stages, including the established interzone-associated genes Gdf5, Sfrp2, and Creb5. Of these 25 genes, 12 also met the temporal enrichment criteria and were retained for the final comparison (Figure 1G, top). We focused on the knee to investigate regulators contributing to the coordinated formation of articular cartilage, ligaments, and menisci. To examine regional differences in interzone gene expression, we compared the knee and ankle at E12.5 and E13.5. Among the 12 candidates, we prioritized genes more highly expressed in the knee at both stages, yielding Gdf5 and Trps1 (Figure 1G, bottom). Because Gdf5 is an established regulator of joint development, we selected Trps1 for further investigation. *Trps1* encodes an atypical GATA-type transcription factor required for growth plate chondrocyte proliferation, survival, and hypertrophic differentiation ^27–29^, for temporomandibular joint development ^30^, and for postnatal hip and patellar morphology ^31^; to our knowledge it has no described role in interzone specification. In the spatial data, *Trps1* was sharply and transiently induced at E12.5, as were *Gdf5*, *Creb5*, and *Sfrp2*; at E11.5 and E12.5 the *Trps1*-positive spot fraction and mean expression exceeded those of *Gdf5*, and from E13.5 *Gdf5* exceeded *Trps1* (Figure 1H).

**Figure 1.**
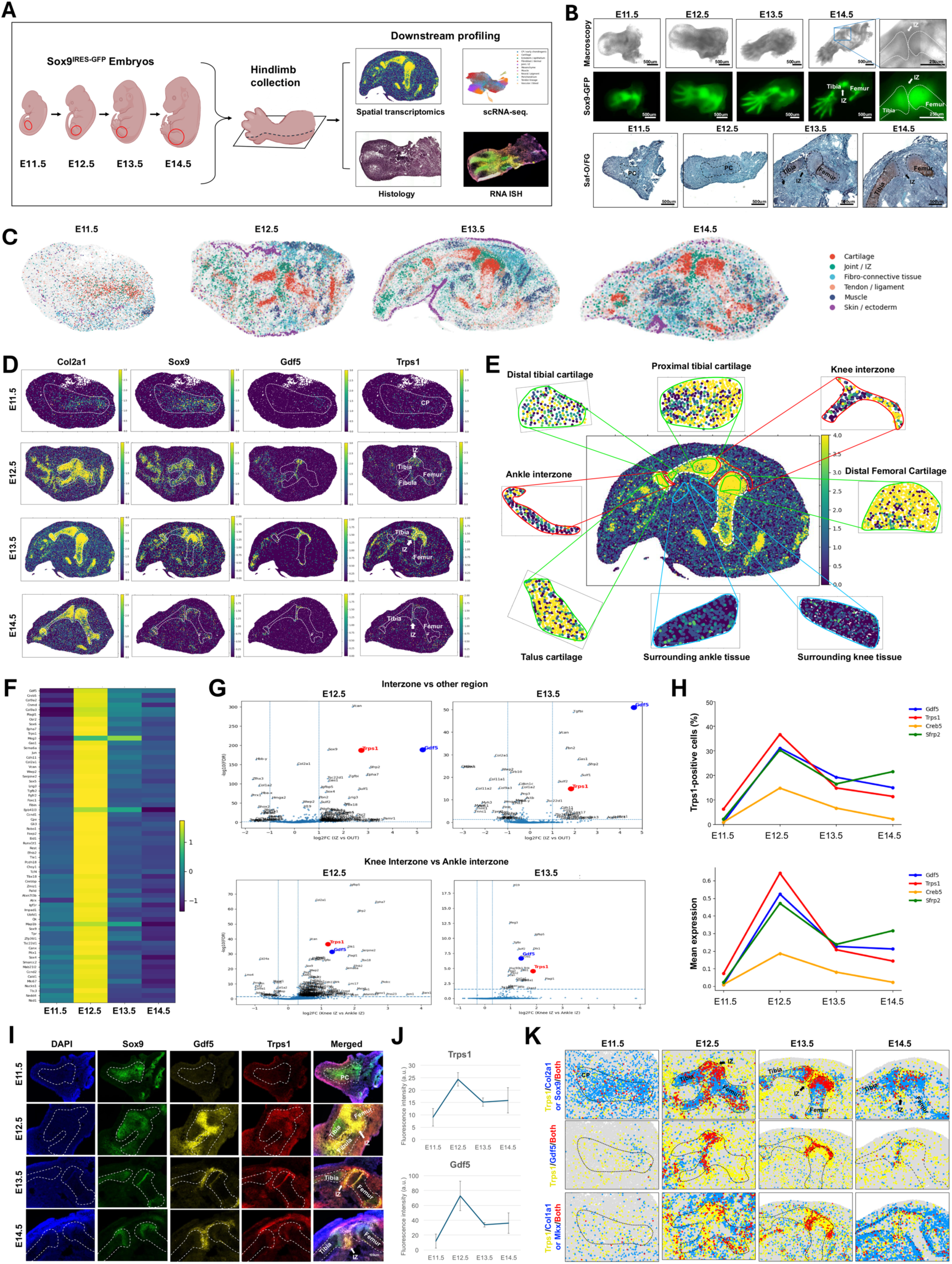
Anatomically matched spatial comparisons nominate *Trps1* as an interzone-enriched candidate regulator. (A) Experimental and analytical workflow. Hindlimbs were collected from *Sox9*-IRES-GFP embryos at embryonic day (E) 11.5, E12.5, E13.5, and E14.5 and analyzed by spatial transcriptomics, single-cell RNA sequencing (scRNA-seq), histology, and RNAscope in situ hybridization (ISH). (B) Brightfield and *Sox9*-GFP images of hindlimbs from E11.5 to E14.5 and Safranin O/Fast Green (Saf-O/FG) sections. Cartilage anlagen condense and a *Sox9*-GFP-low cleft appears between femur and tibia from E13.5. Scale bars as marked. (C) Tissue annotation of the spatial transcriptomic sections at each stage: cartilage, joint/interzone (IZ), fibro-connective tissue, tendon/ligament, muscle, skin/ectoderm. Marker genes used for annotation are listed in Figure S1. (D) Spatial expression of *Col2a1*, *Sox9*, *Gdf5*, and *Trps1* from E11.5 to E14.5. *Trps1* appears in the joint region at E12.5 and extends into the adjacent epiphyseal cartilage. CP, cartilage primordium. (E) Regions annotated for the candidate screen at E13.5, shown on the *Sox9* expression map: knee interzone, ankle interzone, femur distal cartilage, tibia proximal and distal cartilage, talus cartilage, and the surrounding non-cartilage tissue outside the knee and ankle. Regions were drawn by anatomical position and *Col2a1*/*Sox9* expression. (F) The 66 genes enriched in the interzone at E12.5 and E13.5 relative to E11.5 and E14.5. Mean expression across the E11.5 chondrogenic progenitor compartment and the E12.5 to E14.5 interzone regions, row-scaled. (G) Top: differential expression between interzone and surrounding tissue at E12.5 and E13.5 (log2 fold change versus -log10 FDR; Wilcoxon rank-sum with Benjamini-Hochberg correction). Bottom: differential expression between knee and ankle interzones at E12.5 and E13.5. *Gdf5* and *Trps1* are highlighted. (H) Fraction of positive spots (top) and mean expression (bottom) of *Gdf5*, *Trps1*, *Creb5*, and *Sfrp2* in the spatial data by stage. (I) ISH for *Sox9*, *Gdf5*, and *Trps1* with DAPI at E11.5 to E14.5. Dashed lines outline the cartilage anlagen. *Trps1* localizes to the interzone and the adjacent epiphyseal cartilage. Scale bar as marked. (J) ISH fluorescence intensity (arbitrary units) of Trps1 and Gdf5 across developmental stages. Data represent mean ± SD from six embryos per stage. (K) Spatial co-localization of *Trps1* with *Gdf5*, with *Col2a1* or *Sox9*, and with *Col1a1* or *Mkx*, E11.5 to E14.5. Yellow, *Trps1* only; blue, partner only; red, both.

We validated this pattern by multiplexed RNAscope in situ hybridization (ISH). At E11.5, *Trps1* transcript was low in the cartilage primordium. At E12.5, *Trps1* was expressed in the prospective interzone together with *Gdf5*, preceding the morphological cleft at E13.5. At E13.5 and E14.5, *Gdf5* became restricted to the interzone between the tibia and femur, whereas *Trps1* was maintained more broadly across joint-associated structures, including the articular surface and epiphyseal region of the *Sox9*-positive femur and tibia and the adjacent tendon (Figure 1I). ISH intensity of both *Trps1* and *Gdf5* peaked at E12.5 and declined thereafter (Figure 1J). Spatial co-localization analysis showed that *Trps1*-positive spots overlapped with *Gdf5*-positive interzone spots, with the epiphyseal region of *Col2a1*/*Sox9*-positive cartilage, and with *Col1a1*/*Mkx*-positive tendon and ligament domains from E12.5 to E13.5 (Figure 1K). Taken together, these results nominate *Trps1*, from the regional comparison alone, as a candidate regulator that is induced with the interzone and persists in the adjacent epiphyseal cartilage.

### *Trps1* marks a chondrogenesis-biased interzone-associated population

To determine which populations express *Trps1*, we used the scRNA-seq data of the same atlas across the same E11.5 to E14.5 time course ^26^ and integrated the data into a common embedding annotated by stage and population (Figure 2A, B). After quality control, the dataset contained 115,684 cells: 16,011 at E11.5, 59,153 at E12.5, 24,484 at E13.5, and 16,036 at E14.5. *Trps1* was expressed across the joint/interzone (joint/IZ) cluster and chondrogenic populations, whereas *Gdf5* and *Creb5* were restricted to the interzone (Figure 2A). Joint/IZ and cartilage populations expanded from E12.5 to E14.5 (Figure 2C). *Trps1*-positive cells (normalized expression above zero) were 37.6% of cells at E11.5, 44.3% at E12.5, 24.6% at E13.5, and 28.0% at E14.5 (Figure 2D). To determine whether *Trps1* expression extended beyond our knee-focused samples, we examined published mouse limb datasets. *Trps1*-positive fractions were 42.1% in Kelly hindlimbs³² and 52.2% and 47.9% in Zhang hindlimbs and forelimbs³³, respectively (Figure 2E), confirming its expression in additional embryonic limb samples. Stage-specific fractions for *Trps1*, *Sox9*, *Gdf5*, *Myog*, and *Prrx1* are shown in Figure S11. *Trps1* expression peaked at E12.5 and was highest in the joint/IZ population, followed by cartilage, CP/early chondrogenic, and perichondrium (Figure 2F, G). Within the joint/IZ population, 85.5% of cells were *Trps1*-positive at E12.5, 52.6% at E13.5, and 53.6% at E14.5 (Figure 2G).

**Figure 2.**
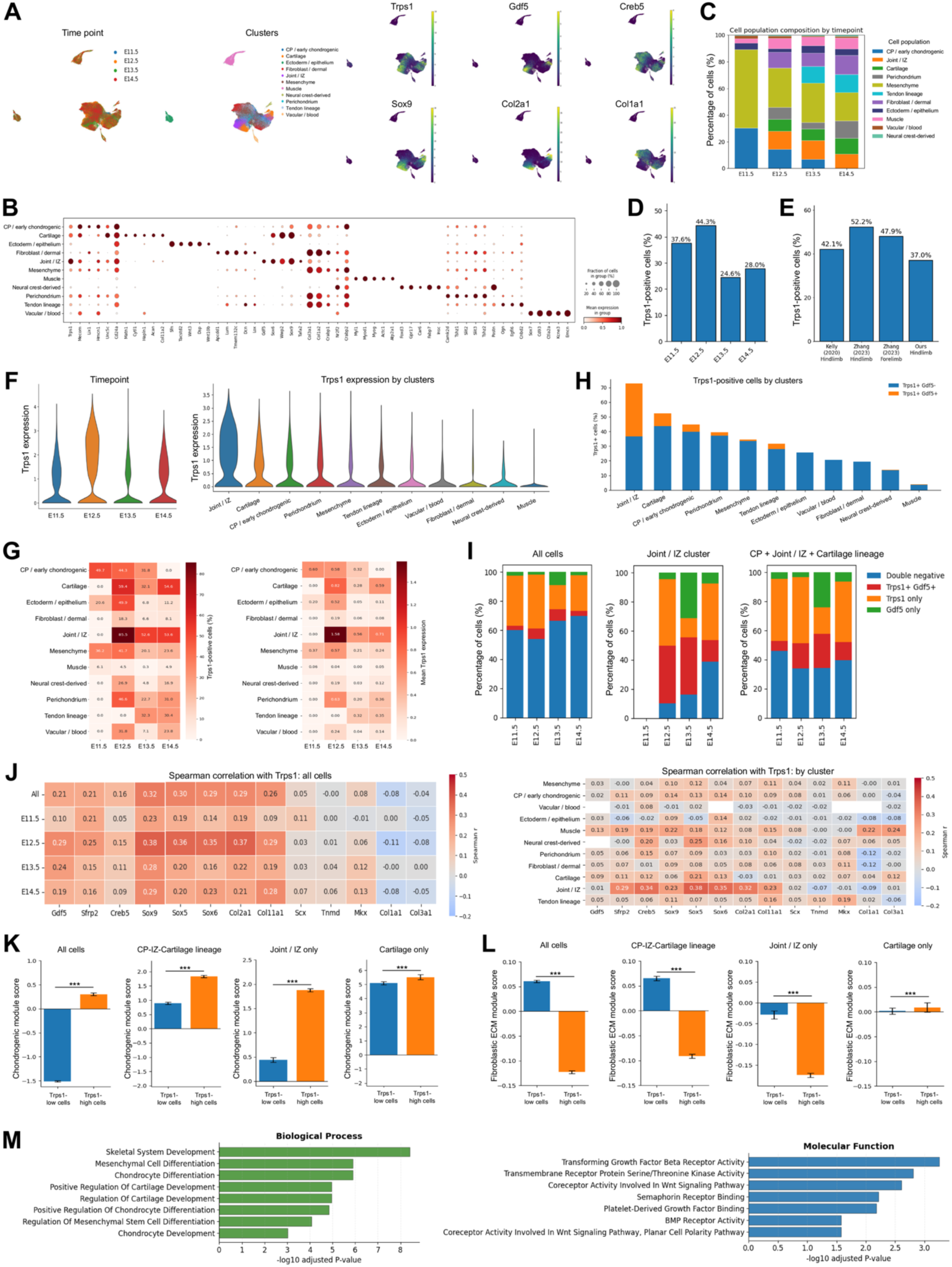
*Trps1* marks a chondrogenically biased interzone-associated population. (A) UMAP of integrated hindlimb scRNA-seq data, E11.5 to E14.5, colored by stage and by population, with feature plots of *Trps1*, *Gdf5*, *Creb5*, *Sox9*, *Col2a1*, and *Col1a1*. (B) Dot plot of markers used to annotate populations. Dot size, fraction of expressing cells; color, mean expression. (C) Population composition by stage. (D) Fraction of *Trps1*-positive cells (normalized expression above zero) by stage: 37.6, 44.3, 24.6, and 28.0%. (E) Overall *Trps1*-positive fraction in the hindlimb data of Kelly et al. ^37^ and (42.1%), the human hindlimb and forelimb data of Zhang, He et al. ^32^ (45.9 and 47.9%), and this study (37.0%). (F) *Trps1* expression by stage (left) and by population (right). (G) *Trps1*-positive fraction (left) and mean *Trps1* expression (right) by population and stage. (H) *Trps1*-positive fraction by population, split into *Trps1*+*Gdf5*- and *Trps1*+*Gdf5*+ cells. (I) *Trps1* and *Gdf5* expression states by stage for all cells, the joint/IZ population, and the combined CP, joint/IZ, and cartilage lineage. (J) Spearman correlation of *Trps1* with interzone, chondrogenic, tendon/ligament, and fibroblastic genes, by stage (left) and by population (right). (K) Chondrogenic module scores in Trps1-low and Trps1-high cells across all cells, the CP/IZ/cartilage lineage, joint/IZ and cartilage populations. (L) Fibroblastic extracellular matrix (ECM) module scores in the same populations. (M) Gene Ontology terms enriched among genes upregulated in *Trps1*-high cells: biological process (left) and molecular function (right), -log10 adjusted P (Enrichr via GSEApy). For (K–L), bars show mean ± SEM; two-sided Mann–Whitney U tests at the single-cell level, ***P < 0.001.

We next asked how *Trps1* expression relates to *Gdf5*. *Trps1*+*Gdf5*+ double-positive cells were concentrated in the joint/IZ population (Figure 2H), where they accounted for 39.52%, 39.25%, and 14.69% of cells at E12.5, E13.5, and E14.5, respectively (Figure 2I). At E12.5, 90.25% of *Gdf5*-positive cells co-expressed *Trps1*, and across all cells and within the combined CP, joint/IZ and cartilage lineage, *Trps1*-only cells outnumbered double-positive cells at every stage (Figure 2I). Thus *Trps1* marks the *Gdf5*-positive interzone and a wider set of joint-lineage and chondrogenic cells. Consistent with this, *Trps1* correlated positively with the interzone genes *Gdf5*, *Sfrp2*, and *Creb5* (Spearman r = 0.21, 0.21, and 0.16 across all cells) and with the chondrogenic genes *Sox9*, *Sox5*, *Sox6*, *Col2a1*, and *Col11a1* (r = 0.32, 0.30, 0.29, 0.29, and 0.26), and showed weak or no correlation with the tendon and ligament genes *Scx*, *Tnmd*, and *Mkx* (r = 0.05, 0.00, and 0.08) or the fibroblastic genes *Col1a1* and *Col3a1* (r = -0.08 and -0.04) (Figure 2J). The correlations were strongest at E12.5 and within the joint/IZ population (r = 0.38 with *Sox5*).

We reasoned that if *Trps1* marks cells biased toward cartilage, *Trps1*-high cells should score higher on a chondrogenic program and lower on a fibroblastic one. We scored each cell for a chondrogenic module and a fibroblastic extracellular matrix (ECM) module and compared Trps1-high and Trps1-low cells. Mean chondrogenic scores were higher in Trps1-high cells across all cells (0.30 versus −1.51), within the CP/joint/cartilage lineage (1.83 versus 0.89), within joint/IZ (1.88 versus 0.44), and, to a lesser extent, within cartilage (5.52 versus 5.09) (Figure 2K). Mean fibroblastic ECM scores were lower in Trps1-high cells across all cells (−0.12 versus 0.06), within the lineage (−0.09 versus 0.07), and within joint/IZ (−0.17 versus −0.03), whereas scores were near zero in both groups within cartilage (Figure 2L). Gene Ontology analysis of genes upregulated in *Trps1*-high cells returned skeletal system development, mesenchymal and chondrocyte differentiation, and cartilage development among biological processes, and TGF-β receptor, Wnt coreceptor, and BMP receptor activity among molecular functions (Figure 2M). Taken together, these data indicate that *Trps1* marks interzone-associated cells biased toward a chondrogenic and away from a fibroblastic program, before matrix differentiation.

### *Trps1*-high interzone cells occupy a transitional state between chondrogenic progenitors and cartilage

To resolve the relationship between *Trps1*-high cells and the chondrogenic lineage, we reclustered the CP/early chondrogenic, joint/IZ, and cartilage cells. This yielded nine populations: Meis/Pbx proximal CP-like progenitors, cycling Meis-positive proximal progenitors, *Ebf1*-positive CP-like progenitors, cycling progenitors, cycling chondrogenic cells, housekeeping/stress-high progenitors, joint/IZ *Trps1*-high cells, a *Cd9*/*Tsc22d1*-positive transition state, and cartilage matrix chondrocytes (Figure 3A and Figure S3). The *Trps1*-high population expressed the interzone genes *Gdf5*, *Creb5*, and *Osr2* and the transition-state genes, and expressed lower levels both of the early CP and proximal markers (*Ebf1*, *Meis1*, *Pbx1*) that define the upstream progenitors and of the matrix genes (*Col2a1*, *Col9a1*, *Acan*, *Matn1*) that define chondrocytes, so it is distinct from both (Figure 3B). Spearman correlation of *Trps1* with interzone and chondrogenic genes was again strongest at E12.5 (r = 0.42 with *Sox9*, 0.46 with *Sox5*, and 0.45 with *Col2a1*) (Figure 3C).

**Figure 3.**
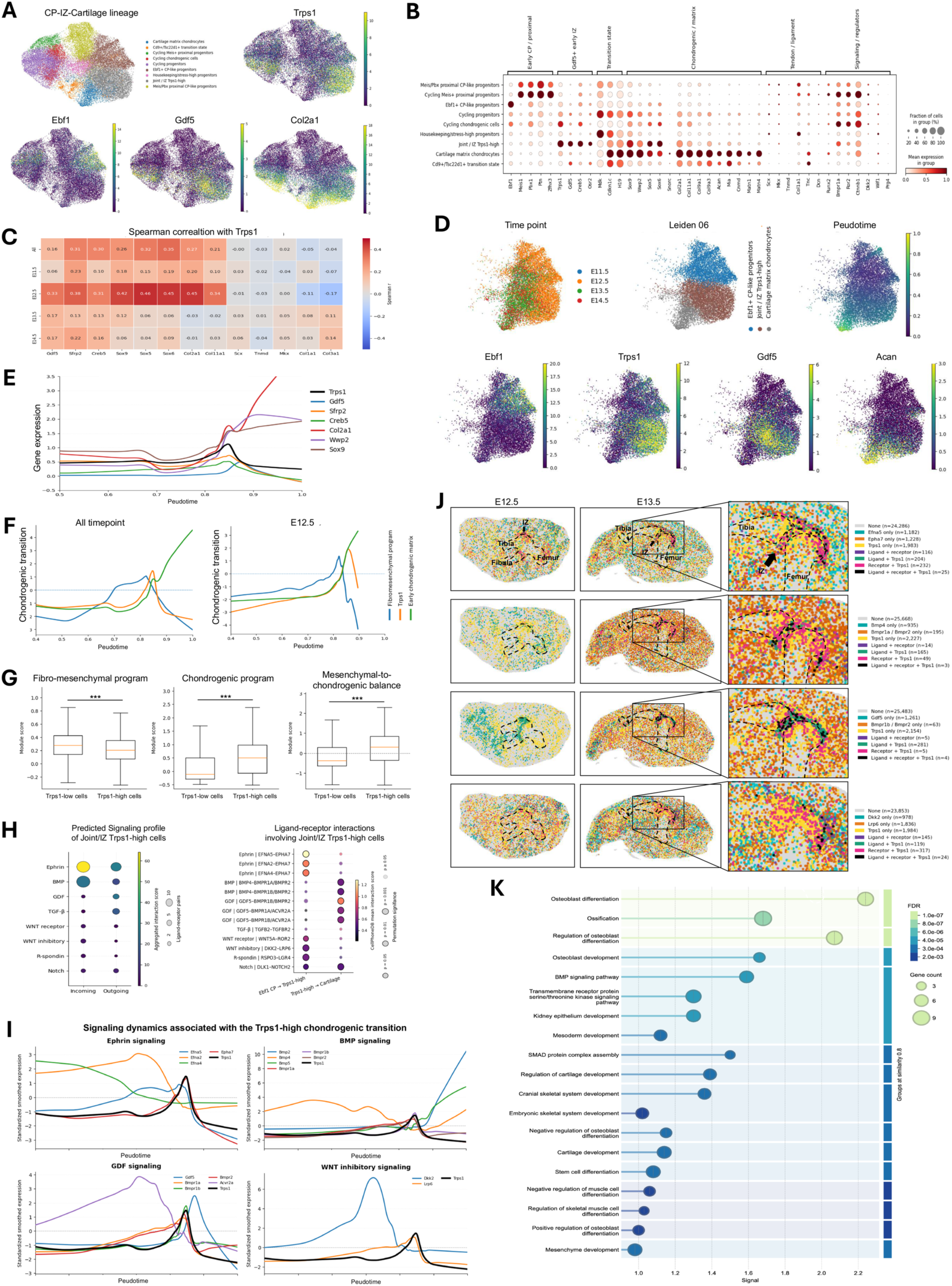
*Trps1*-high interzone cells occupy a transitional state between chondrogenic progenitors and cartilage. (A) Reclustering of CP/early chondrogenic, joint/IZ, and cartilage cells into nine populations, with feature plots of *Trps1*, *Ebf1*, *Gdf5*, and *Col2a1*. (B) Dot plot of genes distinguishing the nine populations, grouped as early CP/proximal, *Gdf5*-positive early IZ, transition state, chondrogenic/matrix, tendon/ligament, and signaling/regulators. (C) Spearman correlation of *Trps1* with interzone, chondrogenic, cartilage matrix, and tendon/fibro-connective genes across all reclustered cells and by stage. (D) The *Ebf1*-positive progenitor, joint/IZ *Trps1*-high, and cartilage matrix chondrocyte populations (Leiden resolution 0.6), colored by stage, population, and diffusion pseudotime rooted in *Ebf1*-positive progenitors, with feature plots of *Ebf1*, *Trps1*, *Gdf5*, and *Acan*. (E) LOWESS-smoothed expression of *Trps1*, *Gdf5*, *Sfrp2*, *Creb5*, *Col2a1*, *Wwp2*, and *Sox9* along pseudotime. (F) Fibro-mesenchymal program, *Trps1*, and early chondrogenic matrix program scores along pseudotime, all stages (left) and E12.5 only (right). (G) Fibro-mesenchymal (left), chondrogenic (middle), and mesenchymal-to-chondrogenic balance (right) scores in *Trps1*-high and *Trps1*-low cells at E12.5. Two-sided Mann– Whitney U tests at the single-cell level; ***P < 0.001. (H) Left: CellChat incoming and outgoing signaling of joint/IZ *Trps1*-high cells. Right: CellPhoneDB ligand-receptor pairs from *Ebf1*-positive CP-like progenitors to *Trps1*-high cells and from *Trps1*-high cells to cartilage matrix chondrocytes; color, mean interaction score; size, permutation P. (I) Standardized LOWESS-smoothed expression of ephrin, BMP, GDF, and Wnt-inhibitory pathway components with *Trps1* along pseudotime. Ephrin components and *Dkk2* peak before or at the rise of *Trps1*; *Bmp4* and *Gdf5* rise during the transition to matrix chondrocytes. (J) Spatial co-expression of *Trps1* with *Efna5*/*Epha7*, *Bmp4*/*Bmpr1a*-*Bmpr2*, *Gdf5*/*Bmpr1b*-*Bmpr2*, and *Dkk2*/*Lrp6* at E12.5 and E13.5, with E13.5 knee enlarged. (K) Functional enrichment of the STRING network generated with Trps1 as the sole input gene, ranked by signal; color, FDR; size, gene count. The network is built from prior protein-interaction knowledge, not from the data in this study.

Diffusion pseudotime rooted in the *Ebf1*-positive progenitors placed the *Trps1*-high population at an intermediate position on a trajectory from progenitors to cartilage matrix chondrocytes (Figure 3D and Figure S4). Along this trajectory, *Trps1* rose together with *Sox9*, *Col2a1*, and *Wwp2* and peaked near pseudotime 0.85, and *Gdf5*, *Sfrp2*, and *Creb5* rose around the *Trps1* peak (Figure 3E). Scoring cells for a fibro-mesenchymal program and an early cartilage matrix program along pseudotime, fibro-mesenchymal expression fell as *Trps1* rose while early cartilage matrix expression rose, both across all stages and at E12.5 alone (Figure 3F). At E12.5, *Trps1*-high cells had lower fibro-mesenchymal scores and higher chondrogenic matrix scores than *Trps1*-low cells (Figure 3G).

To identify signaling associated with this transition, we inferred cell-cell communication among the reclustered populations using CellChat to summarize incoming and outgoing signaling pathway activities (Figure 3H, left) and CellPhoneDB to evaluate individual ligand–receptor interactions, retaining pairs with P < 0.05 (Figure 3H, right). Joint/IZ *Trps1*-high cells were predicted to receive ephrin and BMP signaling, and to send GDF and TGF-β signaling (Figure 3H, left). The ephrin ligands *Efna5*, *Efna2*, and *Efna4*, expressed by *Ebf1*-positive CP-like progenitors, were predicted to bind *Epha7* on *Trps1*-high cells; *Trps1*-high cells expressed *Bmp4* and *Gdf5*, predicted to bind *Bmpr1a*, *Bmpr1b*, *Bmpr2*, and *Acvr2a* on cartilage matrix chondrocytes; and the Wnt inhibitor *Dkk2* was predicted to act through *Lrp6* (Figure 3H, right). Along pseudotime, ephrin ligands and receptors and *Dkk2* peaked before or at the rise of *Trps1*, whereas *Bmp4* and *Gdf5* rose during the transition to matrix chondrocytes (Figure 3I). In the E12.5 and E13.5 spatial data, overlapping expression of Trps1 with ligands or receptors of the Efna5–Epha7, Bmp4–Bmpr1a/Bmpr2, Gdf5–Bmpr1b/Bmpr2, and Dkk2–Lrp6 pairs was observed in the interzone and adjacent epiphyses. At E13.5, ligand–Trps1 overlap was evident within the interzone, while receptor–Trps1 overlap extended into the adjacent epiphyses, with distributions varying among ligand–receptor pairs. Spots co-expressing *Trps1* with both the ligand and the receptor were rare: at E13.5, 25 spots for *Efna5*/*Epha7*, 3 for *Bmp4*/*Bmpr1a*-*Bmpr2*, 4 for *Gdf5*/*Bmpr1b*-*Bmpr2*, and 24 for *Dkk2*/*Lrp6* (Figure 3J). The spatial data are therefore consistent with, but do not establish, signaling between the interzone and the epiphyseal cartilage. For context, the known TRPS1 protein interaction network in STRING, built from *Trps1* as the sole input, is enriched for osteoblast differentiation, ossification, BMP signaling, SMAD complex assembly, and cartilage development; this reflects prior knowledge of TRPS1 rather than a result of these data (Figure 3K). Collectively, these analyses associate the Trps1-high transitional state with predicted ephrin and BMP inputs and GDF and TGF-β outputs along the progenitor-to-chondrocyte trajectory.

### The *Trps1* interzone program is retained in hindlimb explant culture

To test *Trps1* function without generating a conditional mutant, we used ex vivo hindlimb explant culture with localized siRNA delivery ^19–21^. Explant culture preserves the lineages, the spatial arrangement of the limb tissues and the axial patterning genes of the hindlimb for three days, with chondrocyte hypertrophy and tendon maturation lagging their in vivo schedule ^26^. E12.5 *Sox9*-IRES-GFP hindlimbs retained Sox9-GFP in the cartilage condensations and developed a knee joint and digits over the culture period (Figure 4A-C), and the joint region of Day 1 and Day 3 explants contained the joint/IZ, CP/early chondrogenic and cartilage populations of the embryonic hindlimb in their in vivo arrangement, with the femur, tibia and interzone identifiable (Figure 4D-H and Figure S5). We therefore asked whether the *Trps1* program is also retained.

**Figure 4.**
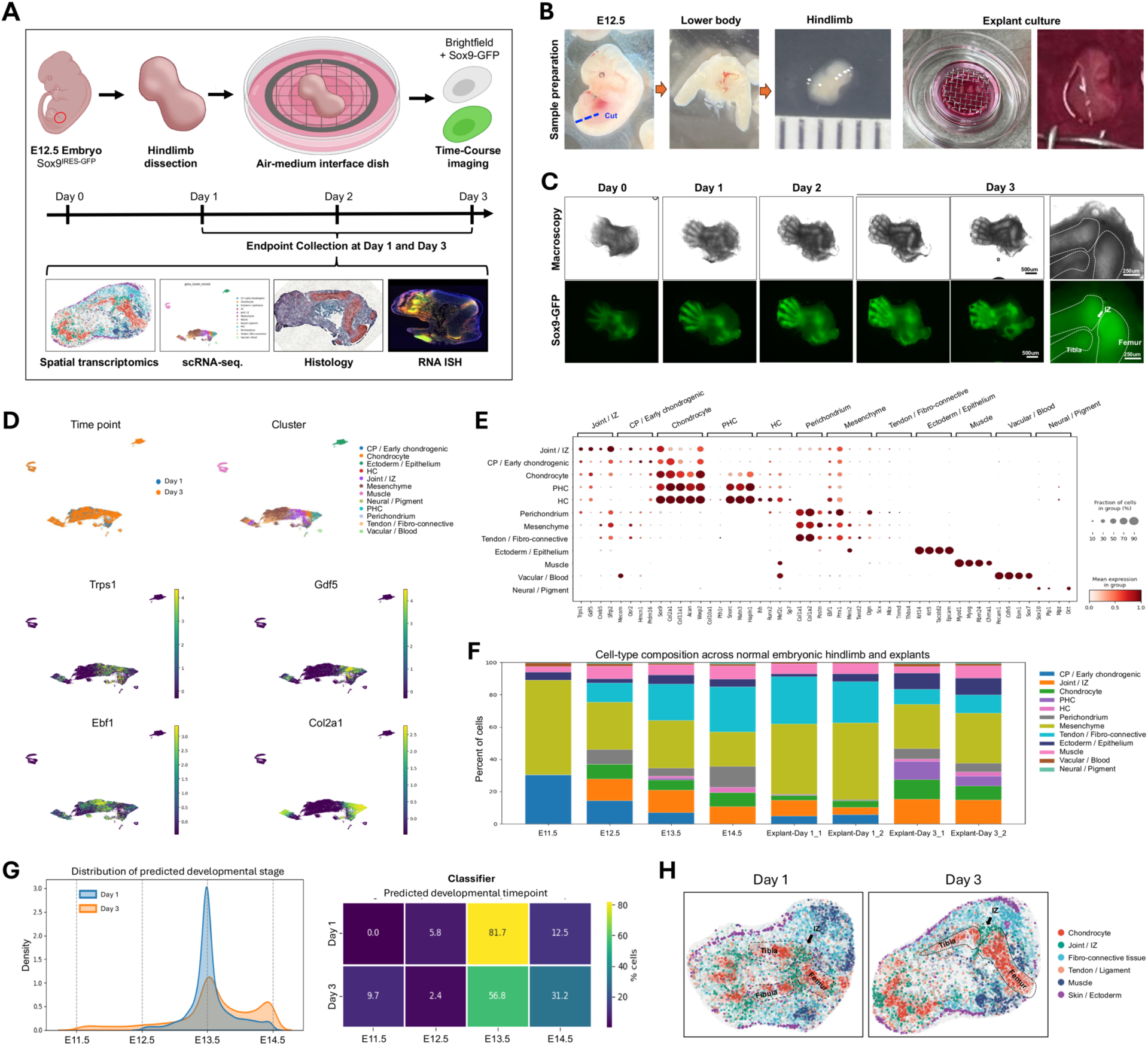
Hindlimb explant culture and the cell populations of the explant joint region. (A) Explant workflow: E12.5 *Sox9*-IRES-GFP hindlimb dissection, air-medium interface culture, time-course imaging, and endpoint scRNA-seq, spatial transcriptomics, histology, and ISH at Day 1 and Day 3. (B) Dissection and culture of an E12.5 hindlimb. (C) Brightfield and *Sox9*-GFP images of explants from Day 0 to Day 3, with the Day 3 knee enlarged. Scale bars as marked. (D) UMAP of Day 1 and Day 3 explant scRNA-seq data colored by day and population, with feature plots of *Trps1*, *Gdf5*, *Ebf1*, and *Col2a1*. (E) Dot plot of markers used to annotate explant populations. PHC, pre-hypertrophic chondrocyte; HC, hypertrophic chondrocyte. (F) Population composition of E11.5 to E14.5 hindlimbs and of two Day 1 and two Day 3 explants. (G) Developmental stage predicted for explant cells by a multinomial logistic regression classifier trained on the E11.5 to E14.5 reference: density of predicted stage (left) and percentage of cells per predicted stage (right). Day 1: 0.0, 5.8, 81.7, 12.5%; Day 3: 9.7, 2.4, 56.8, 31.2% for E11.5, E12.5, E13.5, E14.5. (H) Tissue annotation of spatial transcriptomic sections of Day 1 and Day 3 explants. Femur, tibia, and interzone are outlined.

The *Trps1*-positive fraction rose from 5.77% at Day 1 to 12.53% at Day 3, and mean expression rose in parallel (Figure 5A). Both are lower than in vivo (44.3% at E12.5 and 24.6% at E13.5; Figure 2D) and change in the opposite direction, a difference we attribute to the smaller share of joint and cartilage cells in explants. Within explants, the pattern across populations matched in vivo: joint/IZ had the highest *Trps1*-positive fraction and the highest mean expression, followed by CP/early chondrogenic, perichondrium, and chondrocytes (Figure 5B, C). *Trps1*+*Gdf5*+ double-positive cells were concentrated in joint/IZ, and their share of joint/IZ fell from Day 1 to Day 3 while the *Trps1*-positive share of the cartilage lineage was maintained (Figure 5B, D). *Trps1* correlated with interzone and chondrogenic genes most strongly at Day 1 (r = 0.28 with *Gdf5* and *Sox9*, 0.30 with *Col2a1*) and not with tendon or fibroblastic genes (Figure 5E), as in vivo (Figure 2J). *Trps1-high explant cells had higher mean chondrogenic module scores (0.42 versus −0.04) and lower mean fibroblastic ECM scores (−0.21 versus 0.02) than Trps1-low cells (Figure 5F).* ISH and spatial transcriptomics of Day 1 and Day 3 explants showed *Trps1* and *Gdf5* in the interzone between the *Col2a1*/*Sox9*-positive anlagen (Figure 5G, H), and *Trps1*-positive spots were concentrated at the boundary between cartilage and tendon/fibrous domains within the interzone (Figure 5I). Collectively, the cellular, spatial, and transcriptional features of the *Trps1*-high interzone state were retained in explant culture, at a lower overall *Trps1*-positive fraction.

**Figure 5.**
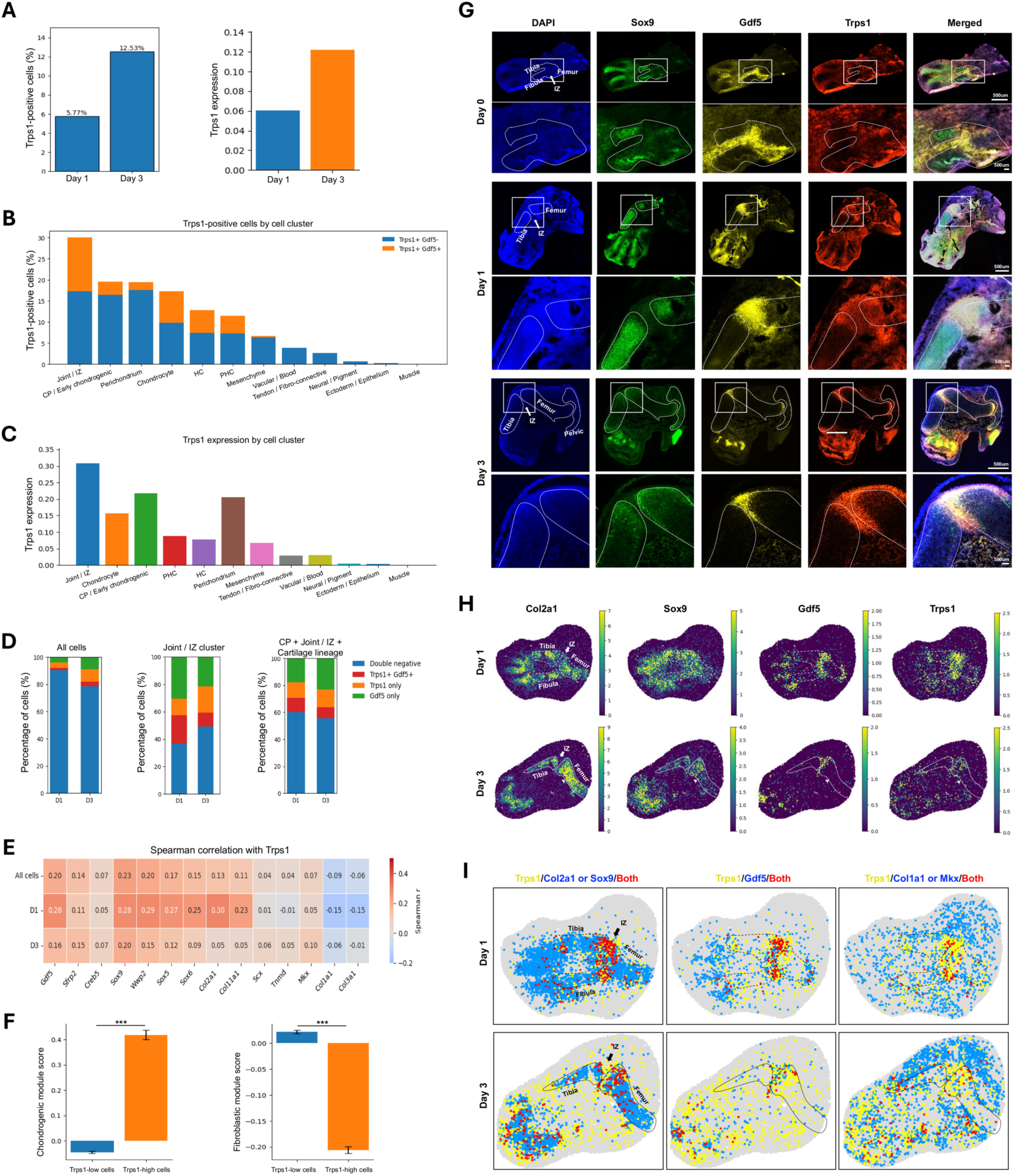
The *Trps1* interzone program is retained in hindlimb explant culture. (A) *Trps1*-positive fraction (5.77 and 12.53%) and mean *Trps1* expression at Day 1 and Day 3. (B) *Trps1*-positive fraction by explant population, split into *Trps1*+*Gdf5*- and *Trps1*+*Gdf5*+ cells. (C) Mean *Trps1* expression by explant population. (D) *Trps1* and *Gdf5* expression states at Day 1 (D1) and Day 3 (D3) for all cells, joint/IZ, and the combined CP, joint/IZ, and cartilage lineage. (E) Spearman correlation of *Trps1* with interzone, chondrogenic, cartilage matrix, and tendon/fibro-connective genes in all explant cells and by day. (F) Chondrogenic and fibroblastic ECM module scores in *Trps1*-low and *Trps1*-high explant cells. (G) ISH for *Sox9*, *Gdf5*, and *Trps1* with DAPI at Day 0, Day 1, and Day 3, with the knee enlarged. Scale bars as marked. (H) Spatial expression of *Col2a1*, *Sox9*, *Gdf5*, and *Trps1* at Day 1 and Day 3. (I) Spatial co-localization of *Trps1* with *Col2a1* or *Sox9*, with *Gdf5*, and with *Col1a1* or *Mkx* at Day 1 and Day 3. Yellow, *Trps1* only; blue, partner only; red, both.

### *Trps1* knockdown in the interzone disrupts joint formation and the flanking cartilage templates

To test whether *Trps1* is required for joint formation, we injected *Trps1* siRNA (assay s211986) complexed with Lipofectamine RNAiMAX into the prospective knee interzone of E12.5 explants at Day 0 and Day 1 and followed development to Day 3 (Figure 6A). A fluorescent control complex delivered the same way remained within the interzone region through Day 3, and *Sox9*-GFP was maintained (Figure 6B). At Day 1, ISH showed reduced Trps1 signal in siTrps1 explants relative to untreated (normal) and negative-control siRNA (siNeg) explants, with Sox9 and Gdf5 largely preserved. Mean Trps1-positive area decreased from 80,275.86 µm² in normal and 79,763.26 µm² in siNeg explants to 36,941.38 µm² in siTrps1 explants, representing reductions of 53.98% and 53.69%, respectively (P = 0.00420 versus normal and P = 0.00463 versus siNeg; Figure 6C–E).

**Figure 6.**
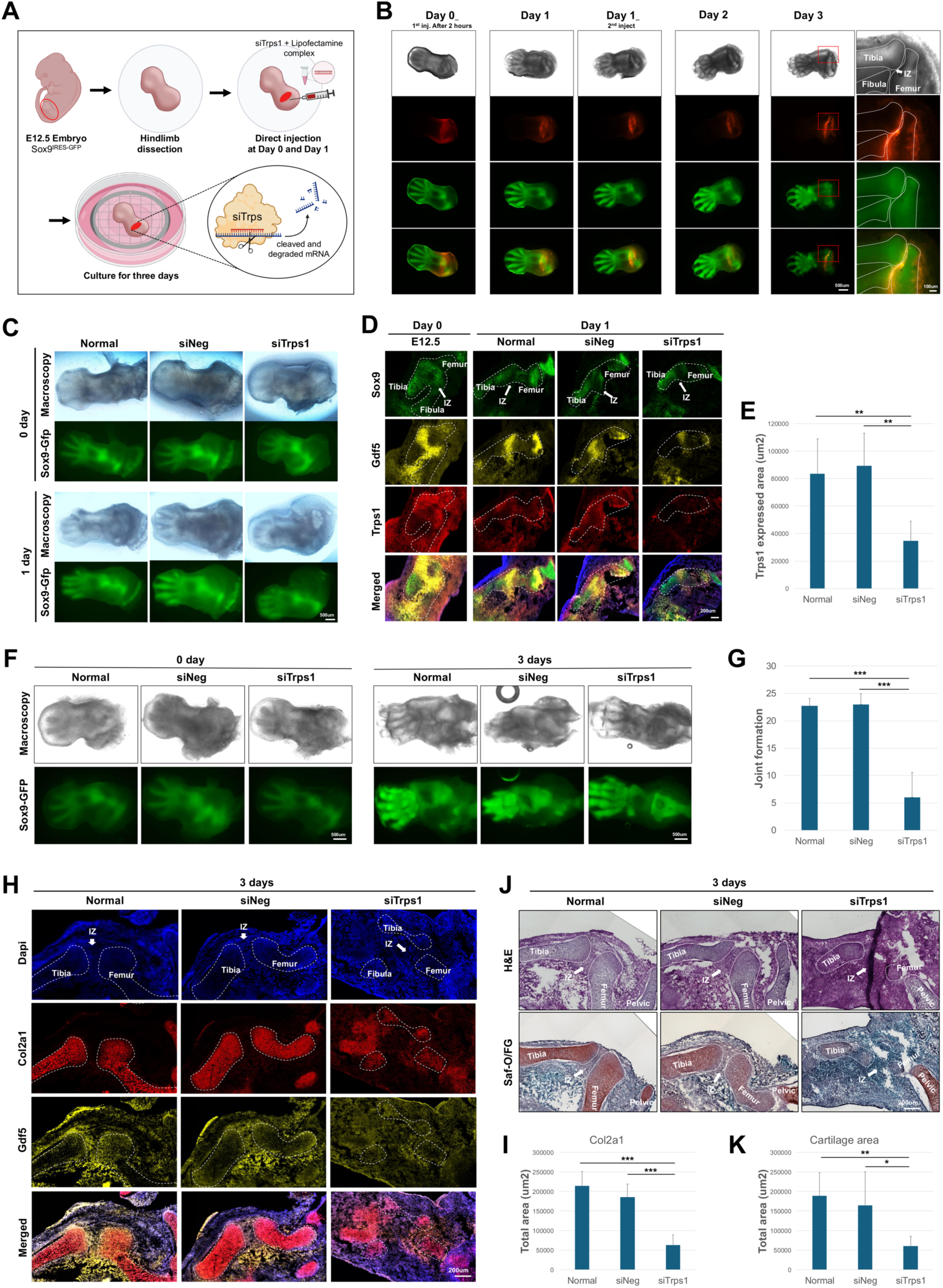
*Trps1* knockdown in the interzone disrupts joint formation and the flanking cartilage templates. (A) Workflow: siRNA-Lipofectamine complex injected into the prospective knee interzone of E12.5 *Sox9*-IRES-GFP explants at Day 0 and Day 1, culture to Day 3. (B) Brightfield, Cy3, and GFP images after injection of the Block-iT Alexa Fluor Red control complex, Day 0 (2 h after the first injection) to Day 3, with the Day 3 knee enlarged. The complex remains within the interzone region. Scale bars as marked. (C) Brightfield and *Sox9*-GFP images of untreated (normal), siNeg, and siTrps1 explants at Day 0 and Day 1. (D) ISH for *Sox9*, *Gdf5*, and *Trps1* at Day 0 (E12.5) and at Day 1 in normal, siNeg, and siTrps1 explants. *Trps1* signal is reduced after siTrps1; *Sox9* and *Gdf5* are preserved. (E) *Trps1*-positive area (um2) at Day 1. (n=7) (F) Brightfield and *Sox9*-GFP images at Day 0 and Day 3. siTrps1 explants do not form separated femoral and tibial templates. (G) Composite joint-formation score at Day 3, the sum of the eight semi-quantitative parameters. (n=7) (H) ISH for *Col2a1* and *Gdf5* with DAPI at Day 3. In siTrps1 explants, *Col2a1* signal is present within the interzone and the remaining signal does not form a template. Dashed lines outline the templates. (I) *Col2a1*-positive area (um2) at Day 3. (n=6). (J) H&E and Saf-O/FG staining at Day 3. (n=6). (K) Cartilage matrix area (um2) at Day 3 from Saf-O/FG sections. For (E, G, I, K), n = 7 explants per group. Data show mean ± SD. Statistical comparisons used one-way ANOVA followed by Tukey’s multiple-comparison test; *P < 0.05, ***P < 0.001.

By Day 3, siTrps1 explants had not formed discrete femoral and tibial templates or a joint between them, whereas normal and siNeg explants had separated joint structures (Figure 6F). A composite joint-formation score, calculated as the sum of eight semi-quantitative parameters (Figure S7), decreased from 22.71 ± 1.38 in normal and 23.00 ± 1.91 in siNeg explants to 6.00 ± 4.55 in siTrps1 explants (P < 0.001 versus either control; Figure 6G). Each of the eight parameters and each of ten quantitative measures (tibial and femoral length, width, area, and integrated density; joint area and integrated density) was reduced in siTrps1 explants (Figure S7, S8). An independent *Trps1* siRNA (assay s211985) produced comparable morphological defects in 6 of 7 explants (Figure S6), which argues against a sequence-specific off-target effect.

ISH at Day 3 showed two changes. In normal and siNeg explants, *Col2a1* was absent from the *Gdf5*-positive interzone and organized into the tibial and femoral templates. In siTrps1 explants, *Col2a1* signal was present within the interzone, and the remaining *Col2a1* signal was diffuse and did not form a template (Figure 6H). Mean Col2a1-positive area decreased from 218,085.67 µm² in normal and 197,962.00 µm² in siNeg explants to 63,622.17 µm² in siTrps1 explants, representing reductions of 70.83% and 67.86%, respectively (P < 0.001 versus either control; Figure 6I). H&E and Saf-O/FG staining showed reduced cartilage matrix and disorganized joint architecture (Figure 6J), with mean cartilage matrix area decreasing from 188,780.00 µm² in normal and 164,678.83 µm² in siNeg explants to 60,290.67 µm² in siTrps1 explants (P = 0.00716 versus normal and P = 0.02723 versus siNeg; Figure 6K). Taken together, *Trps1* knockdown in the interzone prevented joint formation, allowed *Col2a1* expression within the interzone, and reduced formation of the flanking cartilage templates. The template phenotype is consistent with the known requirement for *Trps1* in chondrocytes; the ectopic *Col2a1* is the observation that bears on interzone identity, and we return to the distinction in the Discussion.

### The *TRPS1* program is conserved in the human embryonic knee and redeployed in osteoarthritic cartilage

To determine whether the *Trps1* program is conserved in human joint development, we analyzed published scRNA-seq data of the human embryonic limb at post-conceptional weeks (PCW) 5.1, 5.6, 6.1, 7.2, and 8.4 (1, 6, 6, 2, and 2 embryos), a window comparable to mouse E11.5 to E15.5 ^32^ (Figure 7A-C and Figures S9, S10). The *TRPS1*-positive fraction was 54.1% at PCW5.1, peaked at 63.3% at PCW5.6, and declined to 52.1%, 38.8%, and 38.0% at PCW6.1, 7.2, and 8.4, and mean expression also peaked at PCW5.6 (Figure 7D). *TRPS1* was highest in chondroprogenitors, cycling mesenchyme, and *GDF5*-high chondroprogenitors (Figure 7E). *TRPS1*+*GDF5*+ double-positive cells were most frequent in the chondroprogenitor and *GDF5*-high chondroprogenitor populations, and more than 70% of cells in the combined joint and chondrogenic lineage were *TRPS1*-positive at every stage (Figure 7F, G). *TRPS1* correlated moderately with interzone and chondrogenic genes, most strongly at PCW5.1 and 5.6 (r up to 0.36 with *SOX5*), and not with tendon or fibroblastic genes (Figure 7H). *TRPS1*-high cells had higher mean chondrogenic module scores (0.22 versus −0.13) and lower mean fibroblastic ECM module scores (−0.04 versus 0.03) than *TRPS1*-low cells (P < 0.001; Figure 7I). Spatial transcriptomics of the human knee at PCW5.6, 6.2, and 8.1 showed *TRPS1* in the developing joint, overlapping *GDF5* and *COL2A1*/*SOX9*, with overlap with *MKX*/*COL1A1* in the surrounding tendon and fibro-connective tissue (Figure 7J). Thus, the stage-restricted, chondrogenically biased *TRPS1* program of the mouse interzone is conserved in the human embryonic knee.

**Figure 7.**
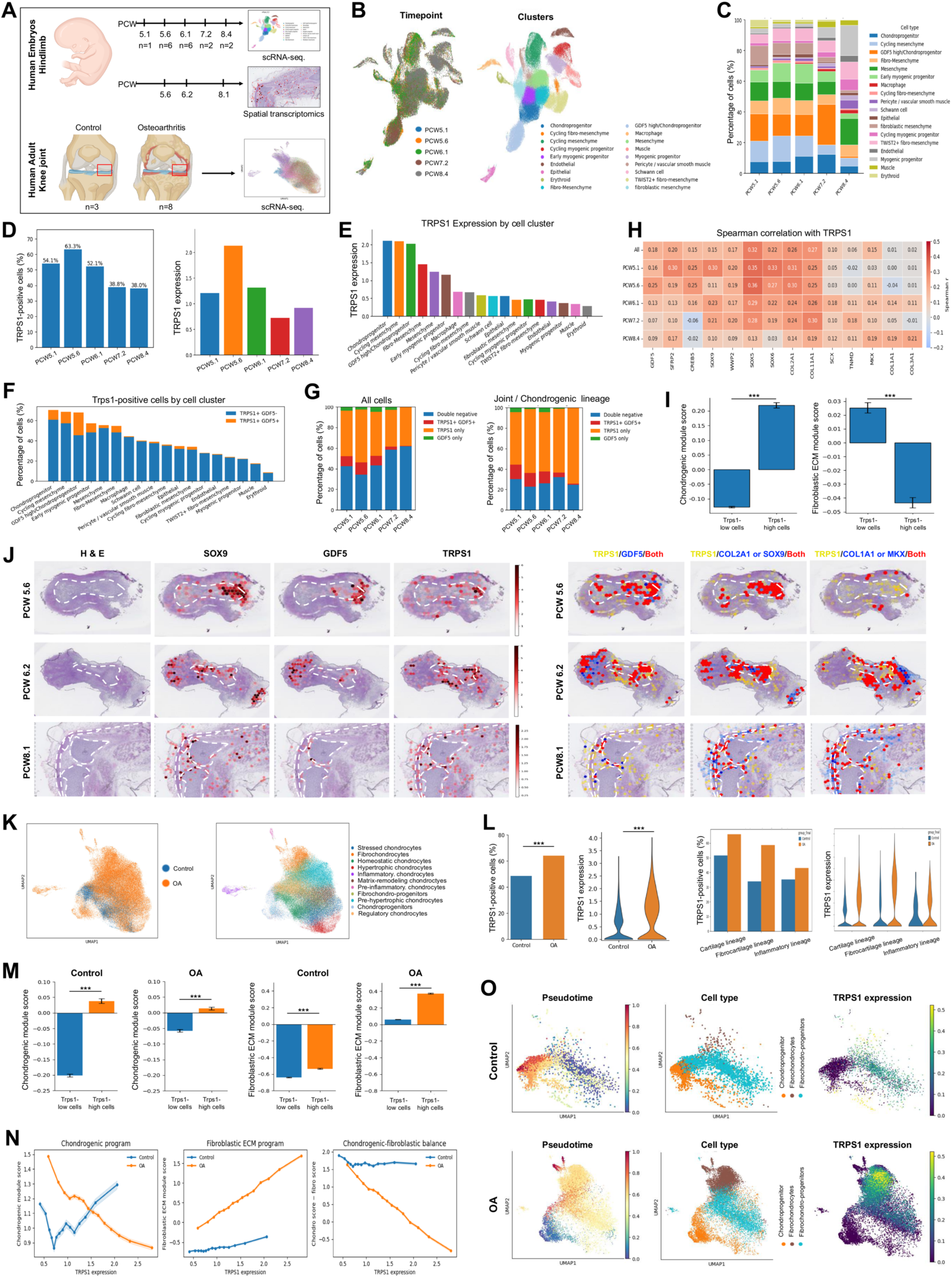
The *TRPS1* program is conserved in the human embryonic knee and redeployed in osteoarthritic cartilage. (A) Human datasets: embryonic hindlimb scRNA-seq at post-conceptional weeks (PCW) 5.1, 5.6, 6.1, 7.2, and 8.4 (n = 1, 6, 6, 2, 2 embryos) and spatial transcriptomics at PCW5.6, 6.2, and 8.1; adult knee articular cartilage scRNA-seq from control (n = 3) and osteoarthritis (OA; n = 8) donors. (B) UMAP of human embryonic limb cells colored by stage and population. (C) Population composition by stage. (D) *TRPS1*-positive fraction (54.1, 63.3, 52.1, 38.8, 38.0%) and mean *TRPS1* expression by stage. Compare the mouse pattern in Figure 2D. (E) Mean *TRPS1* expression by population. (F) *TRPS1*-positive fraction by population, split into *TRPS1*+*GDF5*- and *TRPS1*+*GDF5*+ cells. (G) *TRPS1* and *GDF5* expression states by stage for all cells (left) and the joint/chondrogenic lineage (right). (H) Spearman correlation of *TRPS1* with interzone, chondrogenic, and tendon/ligament genes across all cells and by stage. Compare Figure 2J. (I) Chondrogenic and fibroblastic ECM module scores in *TRPS1*-low and *TRPS1*-high cells. (J) H&E and spatial expression of *SOX9*, *GDF5*, and *TRPS1* in the human knee at PCW5.6, 6.2, and 8.1 (left), and co-localization of *TRPS1* with *GDF5*, with *COL2A1* or *SOX9*, and with *COL1A1* or *MKX* (right). Dashed lines outline the developing joint. (K) UMAP of adult articular chondrocytes from control and OA donors, colored by condition and by chondrocyte state. (L) *TRPS1*-positive fraction and *TRPS1* expression in control and OA cartilage, overall and by cartilage, fibrocartilage, and inflammatory lineage. (M) Chondrogenic and fibroblastic ECM module scores in *TRPS1*-low and *TRPS1*-high cells, control and OA. (N) LOWESS-smoothed chondrogenic score, fibroblastic ECM score, and their difference as a function of *TRPS1* expression in control and OA cells. Shading, 95% confidence interval. (O) Pseudotime, chondrocyte state, and *TRPS1* expression for control (top) and OA (bottom) chondrocytes.

We then asked whether *TRPS1* expression changes in OA. In a published scRNA-seq dataset of articular cartilage from 3 control and 8 OA donors ^25^, restricted to the chondrocyte lineage (Figure 7K and Figure S10), the TRPS1-positive fraction increased from 48.63% in control to 64.00% in OA cartilage, and mean expression increased from 0.48 to 0.97. The largest increase in the positive fraction occurred in the fibrocartilage lineage, from 33.87% to 58.58% (Figure 7L). In control cartilage, *TRPS1*-high cells had higher mean chondrogenic scores than *TRPS1*-low cells (0.04 versus −0.20); this difference was smaller in OA cartilage (0.01 versus −0.06; P < 0.001 for both comparisons). Mean fibroblastic ECM scores were negative in both groups in control cartilage (−0.54 versus −0.64) and positive in OA cartilage, where *TRPS1*-high cells scored higher than *TRPS1*-low cells (0.37 versus 0.06; P < 0.001 for both comparisons; Figure 7M). As a function of TRPS1 expression, chondrogenic scores increased at higher expression levels following an initial decline in control cells, but decreased in OA cells. Fibroblastic ECM scores increased modestly in control cells and more prominently in OA cells, while the chondrogenic–fibroblastic balance remained relatively stable after an initial decline in control cells but decreased markedly in OA cells (Figure 7N). Mapping TRPS1 expression onto separately inferred Palantir trajectories showed elevated expression in OA fibrochondrocytes (Figure 7O). Because the control and OA trajectories were inferred separately, pseudotime values are not comparable between conditions. Taken together, TRPS1 expression is increased in OA cartilage and associated with a fibroblastic matrix program, suggesting context-dependent associations with chondrogenic and fibroblastic programs during development and disease.

## Discussion

When joint progenitors first acquire an identity distinct from the surrounding mesenchyme, and which regulators confer it, are open questions. *Gdf5*-lineage cells enter the interzone over several days ^12^, and transcriptional heterogeneity that correlates with later fate is present at E12.5, before a morphologically distinct interzone exists ^13^. Although single-cell profiling resolves these transcriptional differences, dissociated profiles do not directly retain the anatomical relationships needed to identify genes enriched at a particular joint and developmental stage. Here, we used anatomically matched spatial comparisons of two joints to nominate interzone-enriched genes without a candidate list, and identified *Trps1*, a transcription factor with established roles in the growth plate and in temporomandibular and hip joint development but an unresolved contribution to early knee joint formation^27–30^. *Trps1* marked a chondrogenically biased transitional state at the prospective joint; its knockdown in explants disrupted joint formation and allowed *Col2a1* expression within the interzone; associated expression patterns were observed in the human embryonic knee; and increased TRPS1 expression in human OA cartilage was associated with a fibroblastic matrix program.

First, our spatial screening independently identified a regulator implicated in human skeletal disease. Pathogenic variants in TRPS1 cause tricho-rhino-phalangeal syndrome, characterized by hair, craniofacial, and skeletal abnormalities, including short stature, cone-shaped phalangeal epiphyses, and hip malformations ^33^. Previous studies have demonstrated molecular roles for Trps1 in growth plate chondrocyte proliferation, apoptosis, and differentiation through regulation of Stat3 and interactions with Runx2 and Gli3 ^27–29^. Trps1 loss also disrupts temporomandibular joint development, whereas reduced expression following deletion of intronic regulatory elements causes acetabular dysplasia and other postnatal skeletal abnormalities ^30,31^. However, its spatial and temporal expression in the developing knee between E12.5 and E14.5, and its contribution to interzone and flanking cartilage template formation, have remained unresolved. Our study addresses this gap through expression mapping and localized perturbation. Importantly, Trps1 was nominated through anatomically matched spatial comparisons without using its known disease association, pathway membership, or transcription factor identity as selection criteria. The recovery of a human disease-associated regulator through this unbiased strategy illustrates how spatial comparisons can reveal previously uncharacterized developmental contributions of established disease genes.

Second, the coexistence of ectopic Col2a1 expression within the interzone and impaired formation of the flanking cartilage templates suggests that Trps1 contributes to the spatial organization of chondrogenesis during joint formation. *Trps1* is expressed in the flanking epiphyseal cartilage as well as in the interzone, and the siRNA was delivered into the interzone, so the failure of the femoral and tibial templates may reflect the known requirement for *Trps1* in chondrocytes, loss of a signal from the interzone to the epiphysis, or both. The observation that bears on interzone identity is the *Col2a1* signal within the *Gdf5*-positive domain of siTrps1 explants. The interzone normally excludes chondrogenic fate, and Wnt/β-catenin signaling is required to keep it excluded ^8, 9^. How that positional signal becomes a stable tissue boundary is not understood. Eph receptor and ephrin interactions sort cells between complementary segmental domains to form sharp boundaries ^34^, and our communication and spatial analyses suggest potential signaling relationships among progenitors, the interzone and epiphyseal cartilage. These observations raise the possibility that *Trps1* links positional signaling to the transcriptional states that organize the interzone and adjacent cartilage. However, they do not establish that Trps1 directly represses the chondrogenic program within interzone cells or mediates the predicted ligand–receptor interactions. A conditional deletion in the *Gdf5* lineage and mapping of *Trps1* targets in interzone cells would test this.

Third, the *TRPS1* program is conserved in the human embryonic knee and redeployed in OA. Reactivation of developmental signaling in adult cartilage has been proposed as a mechanism of OA, most notably the hypertrophic program of the growth plate ^22^, and patients with *TRPS1* mutations develop early-onset hip OA ^31^. In the OA atlas, *TRPS1* was increased, concentrated in the fibrocartilage lineage, and associated with a fibroblastic matrix program. Although TRPS1-high cells retained higher chondrogenic scores than TRPS1-low cells, this difference was smaller in OA than in control cartilage. The developmental and OA analyses therefore suggest that the significance of TRPS1 expression depends on the accompanying cellular program, extending the comparison beyond a simple increase in expression. Whether TRPS1 drives the fibrochondrocyte state or marks it cannot be decided from cross-sectional data. Perturbing TRPS1 in OA chondrocytes and measuring both chondrogenic and fibroblastic programs would help distinguish these possibilities.

Direct manipulation of gene activity has long been technically difficult in limb development, and ex vivo whole-limb explant culture offers an alternative that preserves spatial architecture ^19,20,35,36^. Explant culture preserves the lineages, the spatial arrangement of the limb tissues and the axial patterning of the hindlimb for three days ^26^. We found that the *Trps1*-associated transcriptional features of the interzone are retained as well, despite the smaller share of joint and cartilage cells in explants. *Localized siRNA delivery therefore provided a means to test a spatially nominated candidate within its tissue context, and two independent siRNAs supported the morphological phenotype.* This approach enables initial functional testing of spatially identified candidates over a short culture period while preserving the tissue relationships needed to assess joint formation.

We acknowledge that our study is limited in several ways. First, the explant system preserves spatial architecture but not systemic or maternal signals, and its cell composition over-represents fibro-connective and mesenchymal cells. It therefore reproduces selected features of joint development rather than the full in vivo developmental environment. Second, siRNA knockdown is acute and transient and was confirmed at the transcript level only; two independent siRNAs produced the same morphology, and a conditional deletion will be needed to test a chronic requirement and to confine the perturbation to the interzone. Third, *Trps1* is expressed in the epiphyseal cartilage as well as the interzone, so the template phenotype may reflect its chondrocyte function; quantification of *Col2a1* within the *Gdf5* domain and a lineage-restricted deletion will separate the two. Fourth, function was tested in the knee only; the ankle interzone also expresses *Trps1*, at a lower level, and was not tested. Fifth, the role of TRPS1 in human OA was inferred from transcriptomic analyses and requires further functional validation and mechanistic investigation; the OA data are also cross-sectional, from 3 control and 8 OA donors, so pseudotime cannot be read as time and the association between TRPS1 and the fibroblastic program is correlative. Finally, comparisons between *Trps1*-high and *Trps1*-low cells use cells as the unit of test, so their P values overstate certainty; we therefore report effect sizes, and the direction of each effect was reproduced in the explant and human datasets.

Together, our findings identify *Trps1* as a regulator of early knee joint development and suggest that its contribution involves the spatial organization of joint and cartilage formation. Its expression to chondrogenic cell states during development and a fibroblastic matrix program in OA. Defining its direct targets and lineage-specific functions will help explain how these context-dependent associations arise. More broadly, combining anatomically matched spatial comparisons with localized explant perturbation provides a framework for identifying candidate regulators and testing their function within preserved tissue architecture.

## Materials and Methods

### Experimental animals

All animal experiments were approved by the Johns Hopkins University Institutional Animal Care and Use Committee (IACUC; MO23M290) and were conducted in accordance with institutional guidelines and the NIH Guide for the Care and Use of Laboratory Animals. C57BL/6J/*Sox9*-IRES-EGFP mice were maintained under specific pathogen-free conditions with a 12-h light/dark cycle and ad libitum access to food and water. Timed pregnancies were established by 4-h mating, and embryos were collected at E11.5, E12.5, E13.5 and E14.5. Embryos from multiple litters were used throughout the study.

The embryonic and explant single-cell and spatial transcriptomic datasets analyzed here were generated as part of a limb development atlas and are described in full there ^26^; the sample numbers relevant to this study are given below. For embryonic single-cell RNA sequencing, dissected tissues from 6–8 embryos within a single litter were pooled. Spatial transcriptomic analyses included 7, 14, 12, and 10 sections at E11.5, E12.5, E13.5, and E14.5, respectively. For explant analyses, two explants per timepoint were used for single-cell RNA sequencing and two explants per timepoint for spatial transcriptomics. Histological and in situ hybridization analyses used n = 6 embryonic samples per stage and n = 7 explants per condition.

### Embryonic hindlimb collection and sample preparation

Pregnant dams were euthanized by CO₂ asphyxiation followed by cervical dislocation, and embryos were collected into ice-cold phosphate-buffered saline. Hindlimbs were dissected under a stereomicroscope (ZEISS Stemi 508) while preserving overall tissue architecture. Samples were processed immediately for single-cell RNA sequencing, embedded in optimal cutting temperature (OCT) compound for spatial transcriptomics, fixed for histological and RNAscope analyses, or maintained as ex vivo explants.

For histology and RNAscope, hindlimbs were fixed in 4% paraformaldehyde for 6 h at 4 °C, sequentially cryoprotected in 10%, 20% and 30% sucrose, embedded in OCT and stored at −80 °C. For spatial transcriptomics, freshly dissected samples were embedded directly in OCT and snap-frozen in liquid nitrogen without prior fixation. For single-cell RNA sequencing, freshly isolated tissues were processed without freezing.

### Ex vivo limb explant culture and siRNA-mediated knockdown

E12.5 *Sox9*-IRES-EGFP hindlimbs were cultured at the air–liquid interface on 1.0-µm transparent PET membrane inserts (Falcon, Cat. 353102) in BGJb medium (Thermo Fisher Scientific, Cat. 12591038) supplemented with 1% Antibiotic-Antimycotic (Thermo Fisher Scientific, Cat. 15240062). Explants were maintained at 37 °C and 5% CO₂ for up to 3 days with daily medium replacement. Brightfield and GFP fluorescence images were acquired using an EVOS M7000 microscope.

For localization of injected material, Block-iT Alexa Fluor Red Fluorescent Control (Thermo Fisher Scientific, Cat. 14750100) complexed with Lipofectamine was injected into the developing interzone. For functional perturbation, explants received one of two independent Trps1-targeting siRNAs (Thermo Fisher Scientific, Cat. 4390771; Assay IDs s211986 and s211985), non-targeting control siRNA (Cat. 4390843), or no treatment. siRNA stocks (100 pmol/μL) were mixed directly with Lipofectamine RNAiMAX (Thermo Fisher Scientific, Cat. 13778100) at a 1:2 volume ratio, following the previously described^36^. The complexes were delivered into the prospective knee interzone using a fabricated glass micro-needle at Day 0 and Day 1, with an injection volume of 10 μL per explant at each administration (Figure 6A).

Within each litter, embryos were randomly assigned to the normal, siNeg, and siTrps1 conditions, with two embryos contributing explants to each condition. Experiments were performed using three separate litters, with all three conditions represented within each litter. Detailed procedures for micro-needle fabrication and localized siRNA delivery have been described previously ^36^. Explants were harvested at Day 1 to assess knockdown efficiency and at Day 3 for morphological, histological, and in situ hybridization analyses. Joint morphology was evaluated using a previously described semi-quantitative scoring system ^36^ by investigators blinded to experimental condition. The eight parameters (*Sox9* condensation, joint shape, shape of the tibial head, shape of the distal femur, tibial length, tibial thickness, femoral length, femoral thickness) were each scored from 0 to 4, yielding a total score ranging from 0 to 32.

### Histology and RNAscope in situ hybridization

Frozen explant samples were cryosectioned at 10 µm and mounted onto Superfrost Plus slides. Sections were stained with hematoxylin and eosin or Safranin O/Fast Green to assess tissue morphology and cartilage matrix deposition. Multiplexed RNAscope in situ hybridization was performed using the RNAscope Multiplex Fluorescent Assay (Advanced Cell Diagnostics, Cat. 323100) with probes targeting *Col2a1*, *Gdf5*, *Sox9* and *Trps1* according to the manufacturer’s instructions. Images were acquired using an EVOS M7000 microscope. Skeletal element area, cartilage area and *Trps1*-or *Col2a1*-positive area were quantified using ImageJ. Fluorescence thresholds were determined separately for each slide based on its background signal to exclude background fluorescence. Col2a1-positive area was quantified within regions of interest delineating the tibial and femoral cartilage templates.

### Spatial transcriptomics and candidate identification

The spatial transcriptomic data analyzed here were generated for the limb development atlas ^26^ as follows. Fresh-frozen embryonic hindlimbs and explants were profiled using the Takara Bio Curio-Seeker platform. Ten-micrometre sections were mounted onto 10 × 10-mm Curio-Seeker tiles for embryonic samples or 3 × 3-mm tiles for explants. Libraries were prepared according to the manufacturer’s protocol and sequenced on an Illumina NovaSeq X to a target depth of approximately 1-1.5 × 10^9^ reads per tile. Raw sequencing data were processed using the Curio-Seeker pipeline (v3.0.0) and aligned to the Mus musculus GRCm38 reference genome.

Spots with fewer than 15 detected genes were excluded. Additional low-quality spots along tissue edges were removed within each manually defined region of interest using the STUF contour profile function applied to total counts per spot. Predicted genes, CT010467.1, and ribosomal and mitochondrial genes were also excluded.

Spatial datasets were analyzed using Scanpy (v1.9). Anatomical regions were manually annotated in CellXGene (v1.3.0) using tissue morphology together with *Sox9*, *Col2a1* and *Gdf5* expression. Cartilage anlagen, interzone and surrounding non-cartilage regions were annotated independently in the knee and ankle and used for downstream spatial comparisons.

Candidate interzone-associated genes were identified using sequential filtering based on expression within interzone regions, temporal enrichment during E12.5–E13.5 and differential expression relative to neighboring non-interzone tissue. Differential expression was assessed using the Wilcoxon rank-sum test implemented in Scanpy with Benjamini–Hochberg correction. Genes consistently enriched across developmental stages and anatomical replicates were subsequently compared between knee and ankle interzones to identify regionally enriched candidates.

### Single-cell RNA sequencing and downstream analysis

The single-cell data analyzed here were likewise generated for the atlas ^26^. Single-cell suspensions were filtered through a 100-µm nylon mesh, and TER119/CD45-positive hematopoietic cells were depleted using the EasySep Mouse Mesenchymal Stem/Progenitor Cell Enrichment Kit. Libraries were generated using Illumina PIP-seq V4 chemistry and sequenced to approximately 50,000 reads per cell. Count matrices were processed independently for each developmental stage in Scanpy (v1.9).

Cells were filtered according to library complexity and mitochondrial content. Depending on the dataset, cells were required to contain at least 500–750 total UMIs and at least 1,000 detected genes, while cells with >40,000–65,000 total counts were excluded. Dataset-specific mitochondrial filtering bounds were selected within the reported 1–15% range based on the observed distributions. Genes expressed in fewer than 0.1% of cells were excluded. After QC, the embryonic datasets contained 115,684 cells: 16,011 at E11.5, 59,153 at E12.5, 24,484 at E13.5, and 16,036 at E14.5. The explant datasets contained 12,024 cells at Day 1 and 16,624 cells at Day 3 after QC. Ambient RNA contamination was corrected using SoupX (SoupX v1.3.0). SoupX-corrected counts were normalized to 10,000 counts per cell, log-transformed, and the top 2,000 highly variable genes were identified using the Seurat v3 method prior to principal component analysis. Cell clusters were annotated based on canonical lineage markers and cluster-specific differentially expressed genes. Clustering was optimized using pySCN.tl.cluster_alot(), which iteratively evaluates clustering parameters to maximize the number of cluster-specific marker genes. Subclustering was performed where necessary to further resolve cellular heterogeneity. Cluster robustness was assessed using the pySCN random forest classifier trained on 50% of cells per developmental stage (minimum 50 cells per cluster). Clusters with classification precision below 0.5 were merged with the most closely related cluster and reclassified until transcriptionally distinct populations were obtained. For visualization across developmental stages, datasets were integrated using Harmony with developmental stage as the batch variable. For this integration, 4,000 highly variable genes were selected with developmental stage as the batch key. A nearest-neighbor graph was constructed using 30 Harmony-corrected principal components and 30 nearest neighbors, and UMAP was computed with min_dist = 0.35. The stage-specific cell-population annotations were retained after integration.

Cells were defined as *Trps1*-positive or *Gdf5*-positive when normalized expression exceeded zero and were further classified into *Trps1*−*Gdf5*−, *Trps1*+*Gdf5*−, *Trps1*−*Gdf5*+ and *Trps1*+*Gdf5*+ states. For high–low comparisons, cells with Trps1 expression at or above the 75th percentile were classified as Trps1-high, whereas those at or below the 25th percentile were classified as Trps1-low. Zero-expression cells were included when calculating these thresholds and were retained in the low group. All cells tied at either threshold were included, so group sizes were not necessarily exactly 25% of cells. Differential expression was performed using scanpy.tl.rank_genes_groups() with the Wilcoxon rank-sum test and Benjamini–Hochberg correction. Spearman correlations were calculated between *Trps1* and representative lineage-associated genes.

Chondrogenic and Fibroblastic ECM module scores were calculated using scanpy.tl.score_genes() with predefined gene sets (Chondrogenic genes; *Sox9*, *Sox5*, *Sox6*, *Col2a1*, *Col11a1*, *Col9a1*, *Col9a3*, Fibroblastic ECM genes; *Col1a1*, *Col1a2*, *Tnc*, *Dcn*, *Lum*). A chondrogenic–fibroblastic balance score was calculated as the chondrogenic module score minus the fibroblastic ECM module score. Gene Ontology enrichment analysis of genes upregulated in *Trps1*-high cells was performed using Enrichr through GSEApy with the GO Biological Process 2023, Cellular Component 2023, and Molecular Function 2023 libraries.

### Comparison with published mouse limb datasets

To determine whether Trps1 expression was also observed in independent embryonic limb samples, we analyzed published mouse scRNA-seq datasets from Kelly et al.³² and Zhang et al.³³. Analyses included E11.5, E13.5, and E15.5 hindlimbs from the Kelly dataset, and E12.5 and E13.5 hindlimbs and E11, E12, E13, E13.5, and E14 forelimbs from the Zhang dataset. These datasets sampled whole limbs, with the Zhang hindlimb analysis combining proximal, middle, and distal regions, whereas our E12.5–E14.5 samples were harvested specifically from the knee region.

Trps1-positive cells were defined by normalized expression greater than zero, and fractions were calculated across cells pooled within each dataset and limb type. Developmental stages, anatomical sampling, and tissue-processing procedures differed among datasets.

### Joint-lineage reclustering, pseudotime and cell–cell communication analysis

Chondrogenic progenitor, joint/interzone, and cartilage populations were extracted and reclustered using the preprocessing workflow described above, with 4,000 highly variable genes selected for dimensionality reduction. A nearest-neighbor graph was constructed using 20 Harmony-corrected principal components and 30 nearest neighbors, followed by Leiden clustering at a resolution of 0.6.

Developmental trajectories were reconstructed using diffusion pseudotime implemented in Scanpy. Diffusion maps were generated using scanpy.tl.diffmap(), followed by pseudotime inference with scanpy.tl.dpt(). The root population was assigned to early *Ebf1*-positive chondrogenic progenitors. Gene-expression dynamics along pseudotime were visualized using LOWESS smoothing implemented in statsmodels with a smoothing fraction of 0.18 for analyses across all developmental stages and 0.22 for E12.5-restricted analyses.

For the trajectory-associated analyses separate gene sets were used to score fibro-mesenchymal and early chondrogenic programs. The fibro-mesenchymal gene set comprised Prrx1, Col1a1, Col1a2, Col3a1, Col5a1, Col5a2, Dcn, Lum, Postn, Fstl1, Sned1, Vim, Ebf1, Fn1, Tnc, and Sparc. The early chondrogenic gene set comprised Sox9, Sox5, Sox6, Wwp2, Col2a1, Col11a1, Col11a2, Col9a1, Col9a2, and Col9a3. Scores were calculated using scanpy.tl.score_genes() with use_raw=False and random_state=0, retaining genes present in the expression matrix. Neither Trps1 nor Gdf5 was included in these gene sets.

Cell–cell communication was inferred using CellChat (v2.1.2) and independently evaluated using CellPhoneDB (v5.0). Incoming and outgoing pathway activities were calculated using the reclustered joint-lineage populations. CellPhoneDB interactions with P < 0.05 were considered significant. Spatial localization of selected ligand–receptor pairs was subsequently assessed using the embryonic spatial transcriptomic datasets. Expression values used for spatial overlays were percentile-clipped (2nd–98th percentile), min–max scaled and thresholded at 0.15 using custom Python scripts.

Protein–protein interaction (PPI) analysis was performed using STRING (v12.0; https://string-db.org/), with Trps1 as the sole input gene. A minimum required interaction score of 0.700 (high confidence) was applied to construct the network. Functional enrichment results for the resulting TRPS1-associated protein network were obtained from STRING’s Functional enrichment output.

### Analysis of ex vivo transcriptomic datasets

The Day 1 and Day 3 explant datasets were also generated for the atlas ^26^. They were analyzed using the same single-cell and spatial transcriptomic workflows applied to embryonic samples. Spatial transcriptomic analyses used two explants per timepoint. Cell-type composition was compared with the in vivo reference atlas. Developmental-stage probabilities were estimated using a multinomial logistic regression classifier implemented in scikit-learn. Among genes shared by the reference and explant datasets, the 3,000 most variable genes with mean expression >0.01 in the embryonic reference were selected as features. Reference cells were randomly downsampled to 5,000 per stage. A stratified 75:25 split provided 3,750 training cells and 1,250 held-out cells per stage for internal cell-level validation (random_state=0). Features were standardized using StandardScaler. The classifier used L2 regularization (C=1.0), balanced class weights, the lbfgs solver, and a maximum of 3,000 iterations. Overall held-out accuracy was 99.76%, with class-specific recall of 99.68%, 100%, 99.52%, and 99.84% for E11.5, E12.5, E13.5, and E14.5, respectively. The classifier was then refitted using all 20,000 sampled reference cells and applied to explant cells without cell-type annotations as predictors. Trps1 expression, Trps1/Gdf5 co-expression states, lineage-associated correlations, and module scores were calculated using the same definitions applied to the embryonic datasets.

### Human embryonic and osteoarthritis datasets

Publicly available human embryonic limb single-cell RNA-sequencing and spatial transcriptomic datasets spanning PCW5.1–8.4 were obtained from Zhang et al. ^32^. Single-cell datasets were processed in Scanpy and integrated across developmental stages using Harmony. Cell identities were taken from the original study and refined using canonical markers where necessary. *TRPS1* expression, *TRPS1*/*GDF5* co-expression, lineage-associated correlations and module scores were analyzed using the same framework applied to the mouse datasets. Spatial datasets from PCW5.6, PCW6.2 and PCW8.1 were used to examine localization of *TRPS1* and joint-associated markers.

Publicly available adult human articular cartilage single-cell datasets from control and osteoarthritis samples were obtained from Fan et al. ^25^. Data were processed and integrated using Scanpy and Harmony, and analyses were restricted to the chondrocyte lineage. *TRPS1*-positive cell fractions, *TRPS1* expression, and chondrogenic and fibroblastic extracellular matrix (ECM) module scores were compared between control and OA cartilage.

For pseudotime analysis, progenitor-like chondrocytes, pre-fibrochondrocytes and fibrochondrocytes were extracted, yielding 3,554 control cells and 16,126 OA cells. Trajectories were reconstructed separately for each condition using Palantir (v1.4.4). Within each condition, 3,000 highly variable genes were selected for principal component analysis, followed by diffusion-map construction with 30 components and multiscale-space calculation. An initial cell was selected from the progenitor-like chondrocytes by maximizing the sum of standardized SOX9 and COL2A1 expression minus standardized COL1A1 and FN1 expression. This cell was supplied as the early-cell input to Palantir, using 500 waypoints and the default boundary-based start-cell selection rather than fixing the supplied cell as the root. *TRPS1* expression was visualized along Palantir pseudotime. To examine the relationship between *TRPS1* expression and lineage programs, chondrogenic and fibroblastic ECM module scores were plotted as a function of *TRPS1* expression using LOWESS smoothing.

### Statistical analysis

Statistical analyses were performed using Python unless otherwise indicated. Biological replicates corresponded to independent embryos, explants, animals or human samples, as specified in the figure legends. Module-score comparisons between Trps1-high and Trps1-low cells were performed at the single-cell level.

Two-group comparisons used two-tailed Student’s t-tests or Mann–Whitney U tests, as appropriate. Paired comparisons used paired t-tests or Wilcoxon signed-rank tests. Comparisons among three or more groups used one-way ANOVA or Kruskal–Wallis tests followed by the multiple-comparison procedures specified in the figure legends. Expression correlations were assessed using Spearman’s rank correlation. Multiple testing in transcriptomic analyses was corrected using the Benjamini–Hochberg method. P < 0.05 was considered statistically significant, using adjusted P values where multiple-testing correction was applied. Significance is indicated as follows: *P < 0.05, **P < 0.01 and ***P < 0.001. Sample sizes and statistical tests are specified in the corresponding figure legends.

### Data and code availability

The mouse single-cell RNA-sequencing and spatial transcriptomic datasets analyzed in this study were generated as part of the limb development atlas ^26^ and are deposited in GEO under accession numbers GSE336117 and GSE346292, to be released upon publication. Sources of the publicly available human embryonic limb and osteoarthritis datasets are provided in the Methods. Analysis code for this study is available at https://github.com/CahanLab/Trps1

## Acknowledgements

The authors acknowledge funding support from the National Institutes of Health (R35GM124725 to P.C.). S.E.S.O. was supported by a Predoctoral Individual National Research Service Award from the National Institute of Arthritis and Musculoskeletal and Skin Diseases (F31AR084301) and an American Dissertation Fellowship from the American Association of University Women (AAUW). J.H.Y. received the Postdoctoral Research Accelerator Award from Johns Hopkins University. Figures were created using BioRender.com. We thank Kaily K. Young for in situ hybridization staining, Kaiwen Chen for slide imaging and Safranin O staining, and Suhani Dewagan for tissue sectioning and imaging.

## Author contributions

J.H.Y. and P.C. conceived and designed the study. J.H.Y. developed the methodology, performed experiments, conducted formal data analyses, and prepared the figures. S.E.S.O. contributed to methodology development and experimental investigations. H.C. contributed to experimental investigations. J.H.Y. wrote the original manuscript. J.H.Y., S.E.S.O., and P.C. reviewed and edited the manuscript. P.C. supervised the study, provided resources, and acquired funding. Claude Opus 5.5 and Fable 5.1 (Anthropic) were used when writing the manuscript to improve conciseness and clarity. The authors have reviewed the manuscript and take full responsibility for its content.

## Declaration of generative AI and AI-assisted technologies

AI-based tools were used to assist with language revision, proofreading, and code development. Their use did not extend to developing the study concepts or drafting the initial manuscript. The authors retain full responsibility for the manuscript’s content.

## Supplementary figure legends

Figure S1. Tissue annotation of the spatial transcriptomic sections at E11.5, E12.5, E13.5, and E14.5: combined map and per-tissue maps for cartilage, joint/IZ, fibro-connective tissue, tendon/ligament, muscle, and skin/ectoderm, with the marker genes used for each.

Figure S2. Fractions of cells expressing Trps1, Sox9, Gdf5, Myog, and Prrx1 by developmental stage in the Kelly mouse hindlimb and Zhang mouse hindlimb and forelimb datasets.

Figure S3. Reclustered CP/IZ/cartilage cells (Figure 3A) colored by stage, with feature plots of *Creb5*, *Sfrp2*, *Wwp2*, and *Sox9*.

Figure S4. The Figure 3D subset with feature plots of *Col2a1*, *Wwp2*, and *Creb5*.

Figure S5. Tissue annotation of Day 1 and Day 3 explant spatial transcriptomic sections, combined and per tissue, with marker genes.

Figure S6. Brightfield and *Sox9*-GFP images of an explant injected with the second *Trps1* siRNA (assay s211985) at Day 1 and Day 3, medial and lateral views.

Figure S7. The eight semi-quantitative joint-morphology parameters at Day 3 in normal, siNeg, and siTrps1 explants, each scored 0 to 4.

Figure S8. Quantitative measures at Day 3: tibial and femoral length, width, area, and integrated density, and joint area and integrated density.

Figure S9. Dot plot of markers used to annotate the human embryonic limb populations (Figure 7B).

Figure S10. Feature plots of *TRPS1*, *LRIG3*, *GDF5*, *SFRP2*, *CREB5*, *SOX9*, *COL2A1*, *COL11A1*, *WWP2*, *EBF1*, *SCX*, *TNMD*, *COL1A1*, and *COL3A1* on the human embryonic limb UMAP.

Figure S11. Feature plots of *TRPS1*, *SOX9*, *COL2A1*, and *COL1A1* on the adult chondrocyte UMAP (Figure 7K).

