## Supplementary for "Anatomically matched spatial comparisons identify *Trps1* as a regulator of knee interzone formation"

### Supplementary 1

E11.5

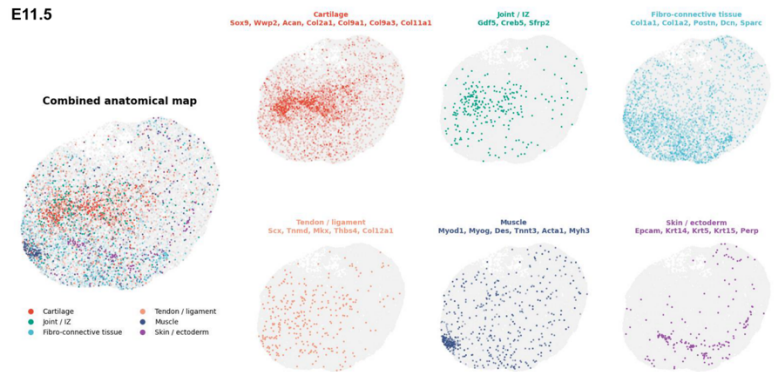

E12.5

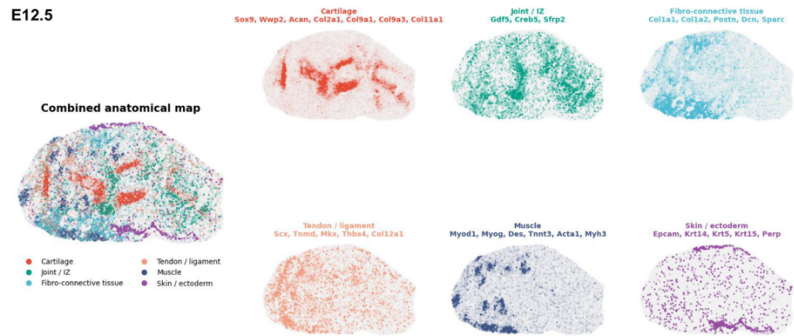

E13.5

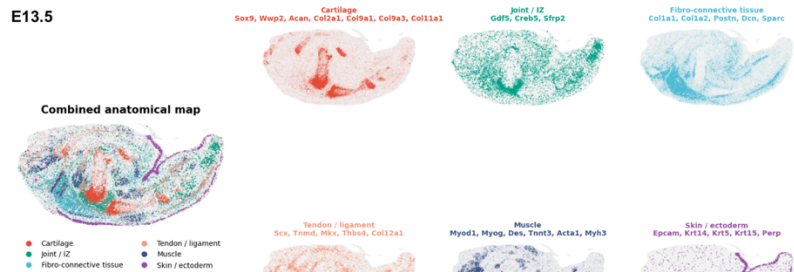

E14.5

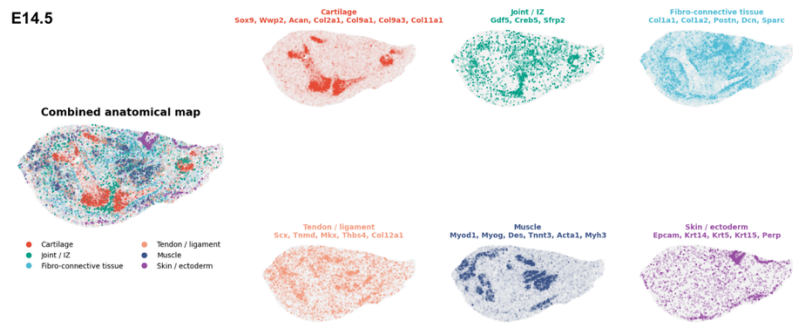

#### Supplementary 2

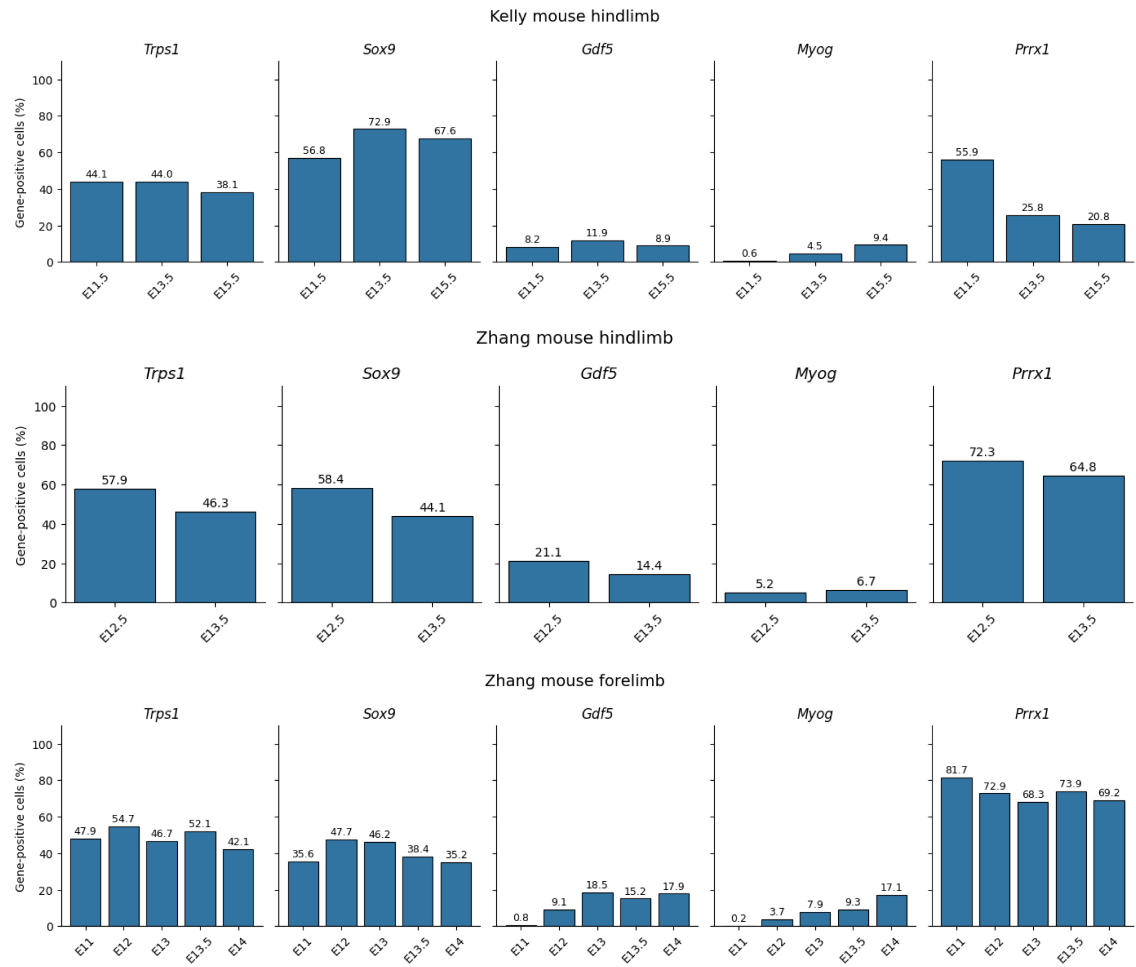

Supplementary 3

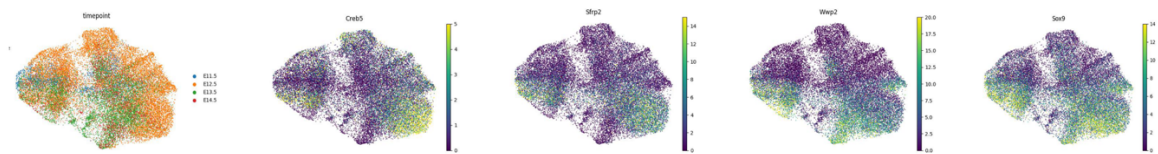

#### Supplementary 4

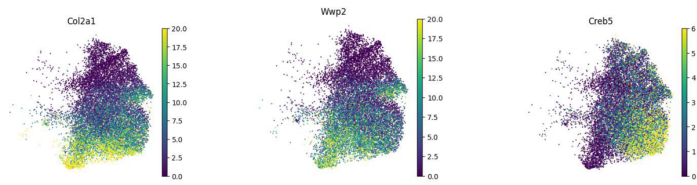

### Supplementary 5

Explant\_Day 1

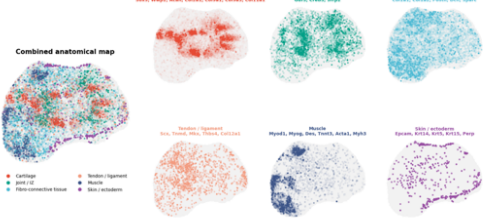

Explant\_Day 3

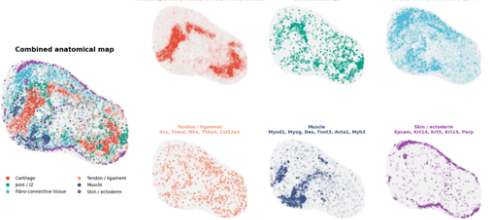

#### Supplementary 6

siTrps1:s211985

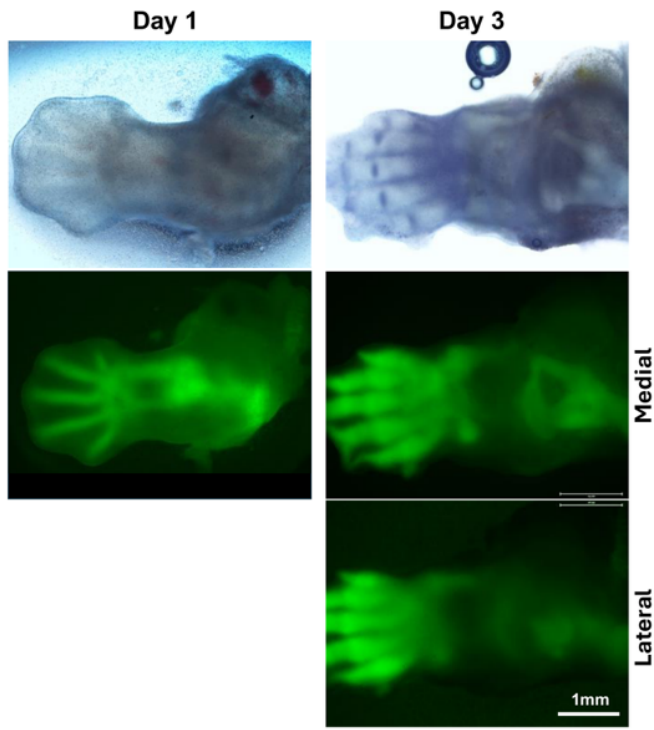

#### Supplementary 7

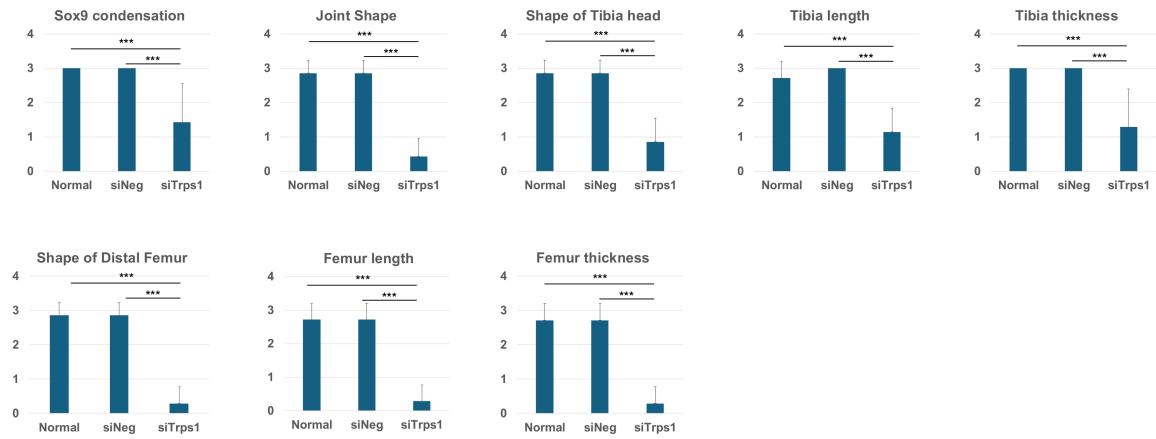

### Supplementary 8

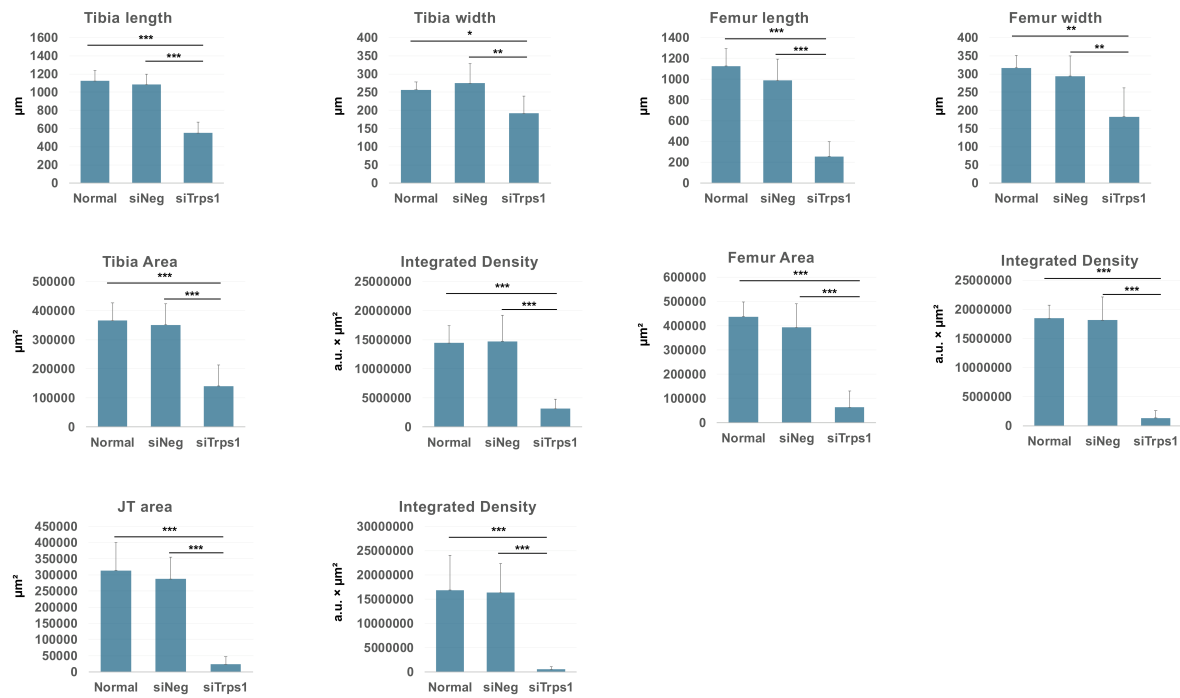

Supplementary 9

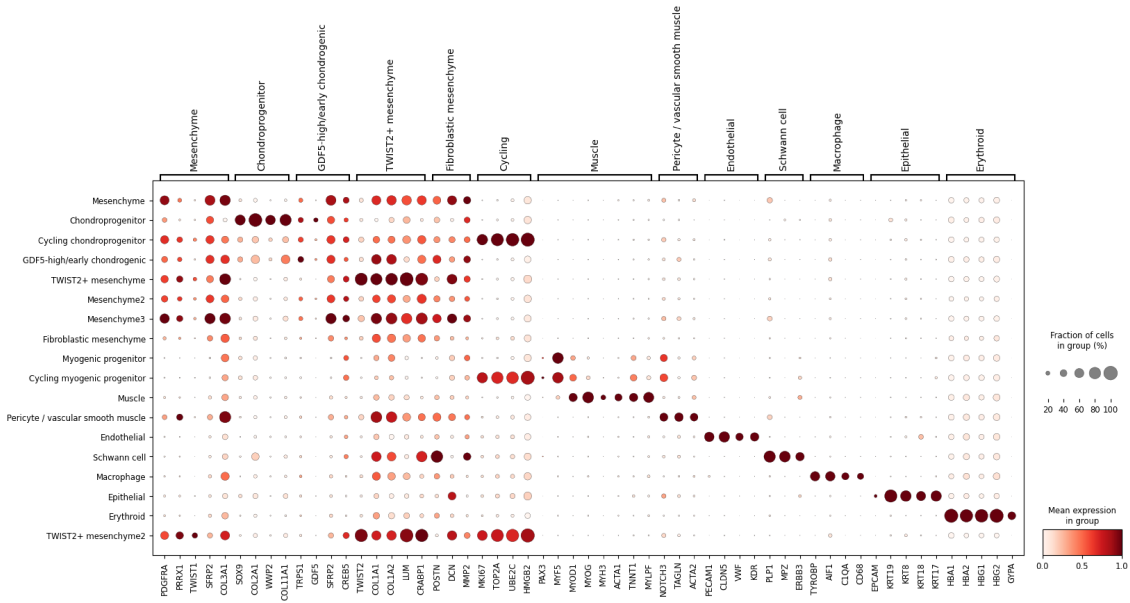

#### Supplementary 10

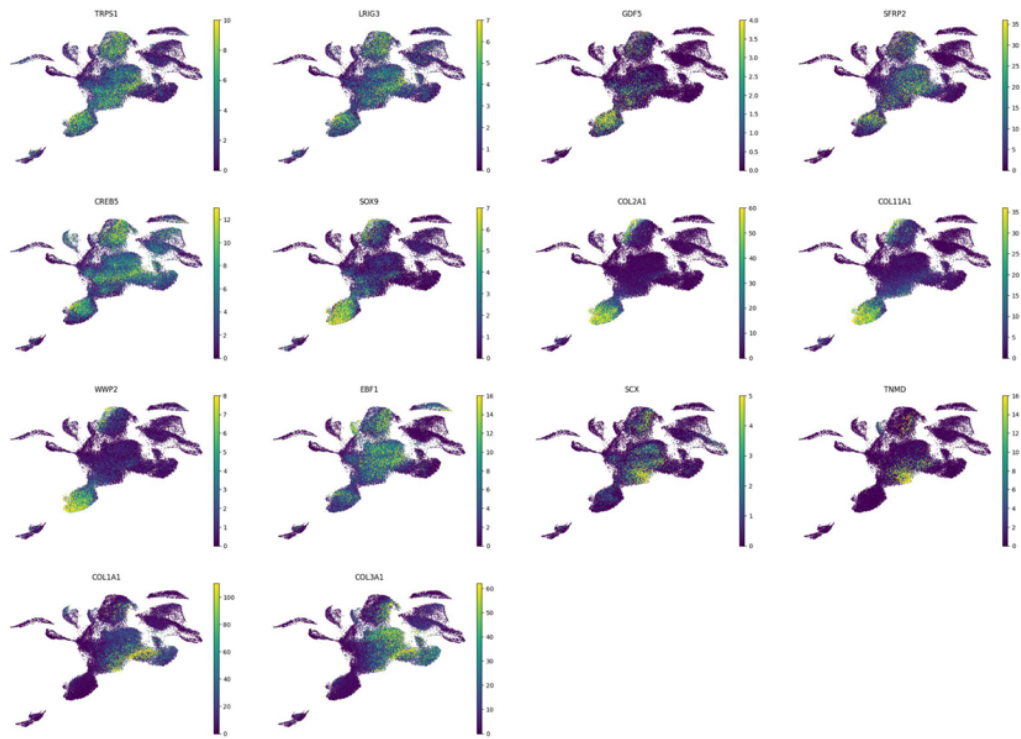

Supplementary 11

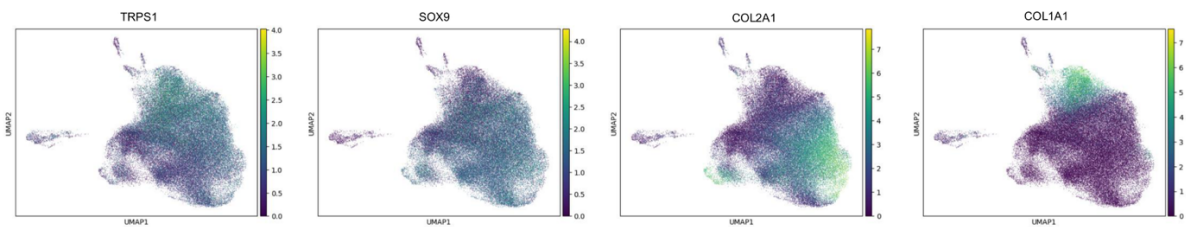
